# Lung biopsy cells transcriptional landscape from COVID-19 patient stratified lung injury in SARS-CoV-2 infection through impaired pulmonary surfactant metabolism

**DOI:** 10.1101/2020.05.07.082297

**Authors:** Abul Bashar Mir Md. Khademul Islam, Md. Abdullah-Al-Kamran Khan

## Abstract

Clinical management of COVID-19 is still complicated due to the lack of therapeutic interventions to reduce the breathing problems, respiratory complications and acute lung injury – which are the major complications of most of the mild to critically affected patients and the molecular mechanisms behind these clinical features are still largely unknown. In this study, we have used the RNA-seq gene expression pattern in the COVID-19 affected lung biopsy cells and compared it with the effects observed in typical cell lines infected with SARS-CoV-2 and SARS-CoV. We performed functional overrepresentation analyses using these differentially expressed genes to signify the processes/pathways which could be deregulated during SARS-CoV-2 infection resulting in the symptomatic impairments observed in COVID-19. Our results showed that the significantly altered processes include inflammatory responses, antiviral cytokine signaling, interferon responses, and interleukin signaling etc. along with downmodulated processes related to lung’s functionality like-responses to hypoxia, lung development, respiratory processes, cholesterol biosynthesis and surfactant metabolism. We also found that the viral protein interacting host’s proteins involved in similar pathways like: respiratory failure, lung diseases, asthma, and hypoxia responses etc., suggesting viral proteins might be deregulating the processes related to acute lung injury/breathing complications in COVID-19 patients. Protein-protein interaction networks of these processes and map of gene expression of deregulated genes revealed that several viral proteins can directly or indirectly modulate the host genes/proteins of those lung related processes along with several host transcription factors and miRNAs. Surfactant proteins and their regulators SPD, SPC, TTF1 etc. which maintains the stability of the pulmonary tissue are found to be downregulated through viral NSP5, NSP12 that could lead to deficient gaseous exchange by the surface films. Mitochondrial dysfunction owing to the aberration of NDUFA10, NDUFAF5, SAMM50 etc. by NSP12; abnormal thrombosis in lungs through atypical PLAT, EGR1 functions by viral ORF8, NSP12; dulled hypoxia responses due to unusual shift in HIF-1 downstream signaling might be the causative elements behind the acute lung injury in COVID-19 patients. Our study put forward a distinct mechanism of probable virus induced lung damage apart from cytokine storm and advocate the need of further research for alternate therapy in this direction.

## Introduction

The recent Coronavirus Disease (COVID-19) pandemic has affected approximately 3.5 million people across 212 countries and territories and the number active cases are still on the rise (at 30 April, 2020) [1]. So far till the writing of this article around 7% of the infected population has suffered death [1] and the fatality rate is continuously increasing due to the lack of detailed knowledge of the molecular mechanism of Severe Acute Respiratory Syndrome Coronavirus 2 (SARS-CoV-2) infection and proper targeted therapeutic approaches against it.

SARS-CoV-2 is a single stranded positive sense enveloped RNA virus and belongs to the betacoronavirus genus of coronavirus [2]. It has 11 protein coding genes encompassing its ~29.9Kb genome [3]. About 90% genomic similarity was observed between SARS-CoV-2 and bat derived SARS-like coronavirus, while SARS-CoV-2 genome is ~79% and ~50% similar with that of Severe Acute Respiratory Syndrome Coronavirus (SARS-CoV) and Middle East Respiratory Syndrome related Coronavirus (MERS-CoV), respectively [2, 4, 5]. Lu et al. (2020) showed the considerable differences between SARS-CoV-2 and SARS-CoV genomes, as in SARS-CoV-2 there has been 380 amino acids substituted, ORF8a deleted, ORF8b elongated, and ORF3b truncated [2]. Though the genomic features are almost similar like those of SARS-CoV, SARS-CoV-2 showed some unique clinical and pathophysiological features like-prolonged incubation period [6], latency inside the host [7] etc. which are making the clinical management of this virus difficult.

Based on the clinical exhibitions of COVID-19, most of the mild to critically affected patients show respiratory complications like- moderate to severe pneumonia, which can further turn into acute respiratory distress syndrome (ARDS), sepsis, and multiple organ dysfunction (MOD) in severely ill patients [8]. Most of these clinical symptoms are associated primarily with respiratory system, specifically with lungs [9] leading to the depleted lung functionality; while the complications with other systems like- cardiovascular system, nervous system etc. were also reported in few patients [10, 11]. Recently cases of pulmonary embolism in the lungs of COVID-19 patients are reported [12].

In SARS-CoV and MERS-CoV infections, increased level of pro-inflammatory cytokines were observed which in turn increased the activation and recruitment of inflammatory cells into the lung tissues, causing the acute lung injury [13]. Similarly, increased levels of many pro-inflammatory cytokines were also detected in moderate to critically affected COVID-19 patients [14], leading to the respiratory failure from ARDS. However the complex interplays between pro-inflammatory and anti-inflammatory cytokines are yet to be completely illustrated. Apart from the cytokine storm, other factors like- host innate immunity, autoimmunity against the pulmonary epithelial and endothelial cells, host genetics and epigenetic factors also play important role in the pathogenesis of SARS-CoV infection [15, 16]. Moreover, the multifaceted host-virus interactions are also found to be a key player in the pathogenesis of previous coronavirus infections [17].

Previously, transcriptional responses in COVID-19 were experimentally recorded using various *in vivo* or *in vitro* approaches like- using cell lines, animal models and COVID-19 infected lungs with SARS-CoV-2 [18], Nasopharyngeal (NP) swab [19], bronchoalveolar lavage fluid of COVID-19 patients [20] etc. which might not provide the actual image of the host transcriptional responses in lung damage upon the viral infection. How the deregulation in the lung’s gene expression profiles is related to the pathogenesis of the SARS-CoV-2 infection and overall pathophysiology of lung failure is not yet devised. Moreover, how the lung’s functionality related pathways are modulated by SARS-CoV-2 is still elusive. As a result, designing of more targeted therapeutic approaches for the clinical management of the COVID-19 patients is still very complicated. In this regard, we have analyzed a publicly available transcriptome data from the lung biopsy of a COVID-19 patient and summarized which pathways are particularly modulated during the SARS-CoV-2 infection, and how the virus is playing a role in the deregulation of the biological processes/pathways related to lungs’ overall functionality and causing acute lung injury in COVID-19 patients.

## Materials and Methods

### Analysis of microarray expression data

Microarray expression data on SARS-CoV infected 2B4 cells or uninfected controls for 24 hrs obtained from Gene Expression Omnibus (GEO), accession: GSE17400 (https://www.ncbi.nlm.nih.gov/geo) [21]. Raw Affymatrix CEL files were background corrected, normalized using Bioconductor package “affy v1.28.1” using ‘rma’ algorithm. Quality of microarray experiment (data not shown) was verified by Bioconductor package “arrayQualityMetrics v3.44.0” [22]. Differentially expressed (DE) between two experimental conditions were called using Bioconductor package Limma [23]. Probe annotations were converted to genes using in-house python script basing the Ensembl gene model (Biomart 99) [24]. The highest absolute expression value was considered for the probes that were annotated to the same gene. We have considered the genes to be differentially expressed, which have FDR [25] p-value ≤ 0.05 and Log2 fold change value ≥ 0.25 (Supplementary file 1).

### Analysis of RNA-seq expression data

Illumina sequenced RNA-seq raw FastQ reads were extracted from GEO database accession: GSE147507 [21]. This data include independent biological triplicates of primary human lung epithelium (NHBE) which were mock treated or infected with SARS-CoV-2 for 24hrs; two technical replicate of post-mortem lung biopsy sample of a COVID-19 deceased patient, along with lung biopsy of two different healthy person. We have checked the raw sequence quality using FastQC program (v0.11.9) [26] and found that the “Per base sequence quality”, and “Per sequence quality scores” were high over threshold for all sequences (data not shown). Mapping of reads was done with TopHat (tophat v2.1.1 with Bowtie v2.4.1) [27]. Short reads were uniquely aligned allowing at best two mismatches to the human reference genome from (GRCh38) as downloaded from USCS database [28]. Sequence matched exactly more than one place with equally quality were discarded to avoid bias [29]. The reads that were not mapped to the genome were utilized to map against the transcriptome (junctions mapping). Ensembl gene model [30] (version 99, as extracted from UCSC) was used for this process. After mapping, we used SubRead package featureCount (v2.21) [31] to calculate absolute read abundance (read count, rc) for each transcript/gene associated to the Ensembl genes. For differential expression (DE) analysis we used DESeq2 (v1.26.0) with R (v3.6.2; 2019-07-05) [32] that uses a model based on the negative binomial distribution. To avoid false positive, we considered only those transcripts where at least 10 reads are annotated in at least one of the samples used in this study and also applied a minimum Log2 fold change of 0.5 for to be differentially apart from adjusted p-value cut-off of ≤ 0.05 by FDR. Raw read counts of this experiment is provided in supplementary file 2. To assess the fidelity of the RNA-seq data used in this study and normalization method applied here, we checked the normalized Log2 expression data quality using R/Bioconductor package “arrayQualityMetrics (v3.44.0)” [22]. From this analyses, in our data no outlier was detected by “Distance between arrays”, “Boxplots”, and “MA plots” methods and replicate samples are clustered together (Supplementary file 3).

### Retrieval of the host proteins that interact with SARS-CoV and SARS-CoV-2

We have obtained the list of human proteins that forms high confidence interactions with SARS-CoV and SARS-CoV-2 proteins from conducted previously studies [33–35] and processed their provided proteins name into the associated HGNC official gene symbol.

### Functional enrichment analysis

We utilized Gitools (v1.8.4) for enrichment analysis and heatmap generation [36]. We have utilized the Gene Ontology Biological Processes (GOBP) [37], Reactome pathway [38], Bioplanet pathways [39], HumanCyc database [40], DisGeNet [41], KEGG pathway [42] modules, and a custom in house built combined module (Supplementary file 4) for the overrepresentation analysis. Resulting p-values were adjusted for multiple testing using the Benjamin and Hochberg’s method of False Discovery Rate (FDR) [25].

### Mapping of the deregulated genes in cellular pathways

We have utilized Reactome pathway browser [38] for the mapping of deregulated genes of SARS-CoV-2 infection in different cellular pathways. We then searched and targeted the pathways which are found to be enriched for lung related functionalities.

### Obtaining the transcription factors which can modulate the differential gene expression

We have obtained the transcription factors (TFs) which bind to the given differentially expressed genes using a custom TFs module created using ENCODE [43], TRRUST [44], and ChEA [45] database.

### Obtaining human miRNAs target genes

We extracted the experimentally validated target genes of human miRNAs from miRTarBase database [46].

### Extraction of transcription factors modulate human miRNA expression

We have downloaded the experimentally validated TFs which bind to miRNA promoters and module it from TransmiR (v2.0) database which provides regulatory relations between TFs and miRNAs [47]. For miRNAs come to play role in transcriptional regulation, we have considered those TFs that are expressed (upregulated) itself and that can ‘activate’ or ‘regulate’ miRNAs, or in absence of TFs (downregulation) that could otherwise ‘suppress’ miRNAs.

### Identification of the host epigenetic factors genes

We used EpiFactors database [48] to find human genes related to epigenetic activity.

### Construction of biological networks

Construction, visualization and analysis of biological networks with differentially expressed genes, their associated transcription factors, associated human miRNAs, and interacting viral proteins were executed in the Cytoscape software (v3.8.0) [49]. We used STRING [50] database to extract highest confidences (0.9) edges only for the protein-protein interactions to reduce any false positive connection.

## Results

### Antiviral immune responses and organ specific functionalities are deregulated in lungs

During a viral respiratory infection, many pathways of the host get fine-tuned to battle with the invading pathogen; on the other side, the infecting viruses also try to hijack and modulate host pathways for immune evasion and survival inside the host [51]. These complex interactions lead to the disease complexity from a viral infection, causing several critical pathophysiological conditions in host respiratory system [51]. To explore which particular biological processes/pathways are deregulated in SARS-CoV-2 infection we first identified the deregulated genes in both SARS-CoV and SARS-CoV-2 infections and performed functional enrichment analyses and then did comparative analyses [36].

We found 3031 (2408 upregulated and 623 downregulated), 142 (91 upregulated and 51 downregulated) and 6714 (2476 upregulated and 4238 downregulated) genes from SARS-CoV infected 2B4 cells, SARS-CoV-2 infected NHBE cells and lung biopsy of COVID-19 patient, respectively (Supplementary Figure 1). We discovered a wide array of genes to be differentially expressed in SARS-CoV-2 infected lung whose expression profiles were observed to be different in SARS-CoV-2 infection in NHBE cells and SARS-CoV infection (Supplementary Figure 1).

As expected, from the enrichment analyses of GOBP, we have observed that biological processes related to antiviral inflammatory responses, viral processes etc. overrepresented in both types of SARS-CoV-2 infections (cell line and primary lung biopsy cells) and SARS-CoV infection (Figure 1A). While several biological processes like- negative regulation of viral replication, immune system process, response to hypoxia, heart development etc. were only enriched for both SARS-CoV-2 infections (Figure 1A), few pivotal processes namely-viral transcription, adaptive immune response, brain development, lung development, respiratory gaseous exchange by respiratory system were uniquely observed for the deregulated genes from COVID-19 affected lung (Figure 1A).

**Figure 1:**
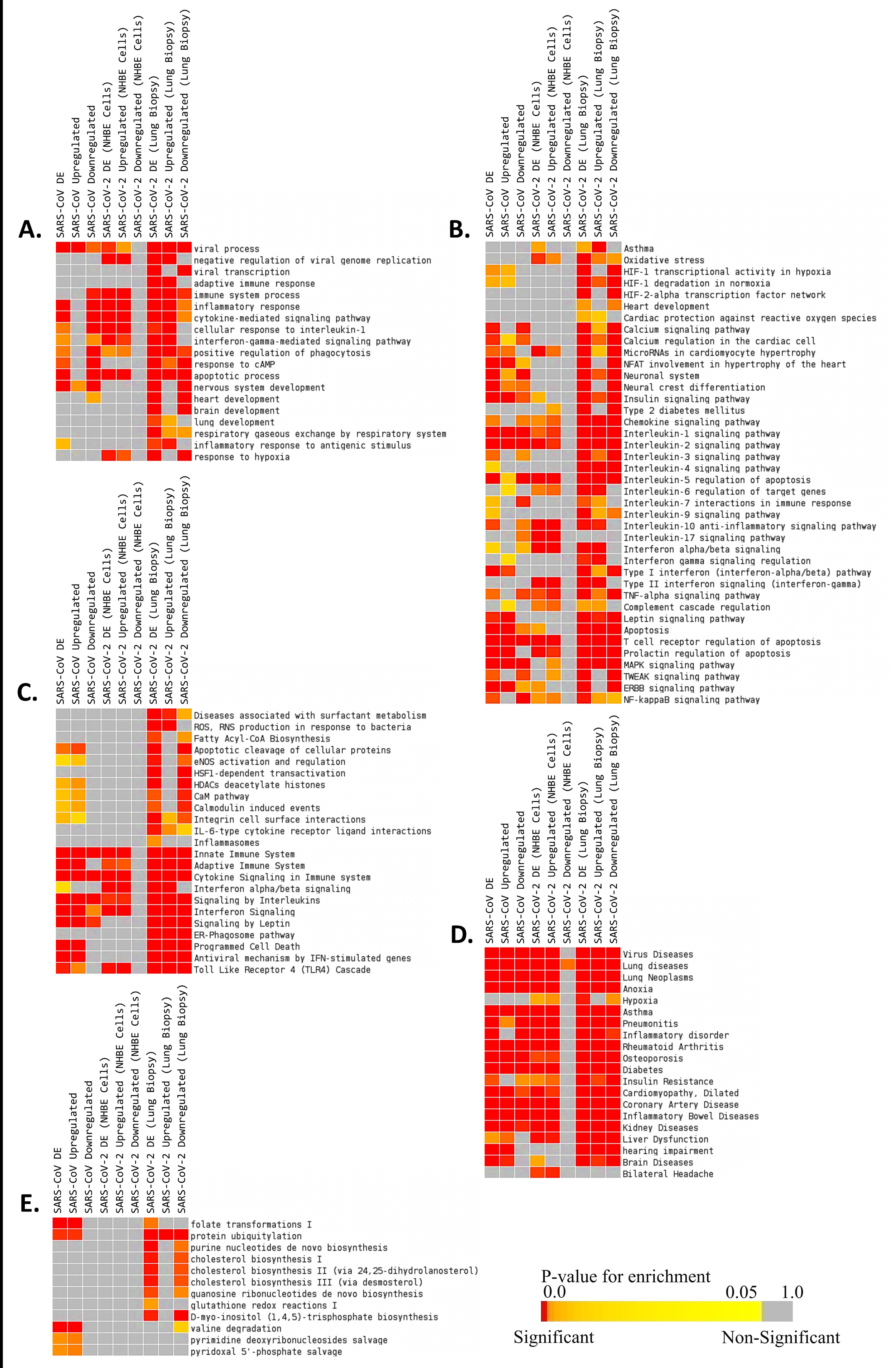
Enrichment analysis and comparison between deregulated genes in SARS-CoV, SARS-CoV-2 (NHBE cells) and SARS-CoV-2 (Lung biopsy) infections using **A.** GOBP module, **B.** Bioplanet pathway module, **C.** Reactome pathway module, **D.** DisGeNet module, **E.** HumanCyc module. Selected significant terms are represented in heatmap. Significance of enrichment in terms of adjusted p-value (< 0.05) is represented in color coded P-value scale for all heatmaps. Color towards red indicates higher significance and color towards yellow indicates less significance, while grey means non-significant. All enriched terms and detail statistics are in Supplementary file 6.

Similarly, from ‘Bioplanet’ module, host antiviral immune responses like- various inflammatory cytokine signaling pathways, apoptosis, interferon-I signaling etc. were activated for all of these infections (Figure 1B, 1C). While, HIF-1 signaling, heart development, asthma, type-II interferon signaling etc. pathways were found in both SARS-CoV-2 infections (Figure 1B, Supplementary Figure 2), curiously, some pathways like-disease associated with surfactant metabolism, ROS/RNS production, fatty acyl-CoA biosynthesis, ER-phagosome pathway, inflammasomes etc. were only found in SARS-CoV-2 infected lung biopsy (Figure 1C, Supplementary figure 2). To further understand, we used the module ‘DisGeNet’ for enrichment analysis which revealed that the deregulated genes of SARS-CoV and SARS-CoV-2 infections are also involved in different diseases like-virus diseases, lung diseases, asthma, pneumonitis, hypoxia etc (Figure 1D, Supplementary figure 2). Interestingly, several cholesterol biosynthesis pathways were found to be deregulated only in the lung biopsy sample of COVID-19 patient (Figure 1E, Supplementary figure 2). As cholesterol in lung plays important roles in maintaining normal lung physiology and protection against many diseases [52], downregulation of these indicates possible association with lung-related comorbidities of COVID-19 patients.

From these enrichment analyses, it is clearly understood that several pathways related to lung’s overall functionality are being deregulated by the infecting SARS-CoV-2 virus. While hunting for more definitive clues on which particular processes are being regulated during this infection, we again performed another round of enrichment analysis with our in house combined module (Supplementary file 4) which gather related modules with few genes to a parent term/process which otherwise left out from analysis due to statistical stringency cutoff (module with 10 genes are selectedly) during enrichment analysis. Surprisingly from this enrichment analysis, we have detected several important lung’s overall functionality related processes only for the deregulated genes from the lung of COVID-19 patient, namely- lung development, pulmonary surfactant metabolism disease or dysfunction, respiratory processes, regulation of respiratory gaseous exchange etc. along with some antiviral responses (Figure 2, Supplementary figure 3). We have also exported enriched genes list of selected significantly enriched terms in color coded heatmaps. Intriguingly, while equating the expression of these enriched genes, we discovered that the genes of these key lung related processes are significantly altered in SARS-CoV-2 infected lungs compared to the SARS-CoV-2 infected NHBE cells and SARS-CoV infection (Supplementary figure 4).

**Figure 2:**
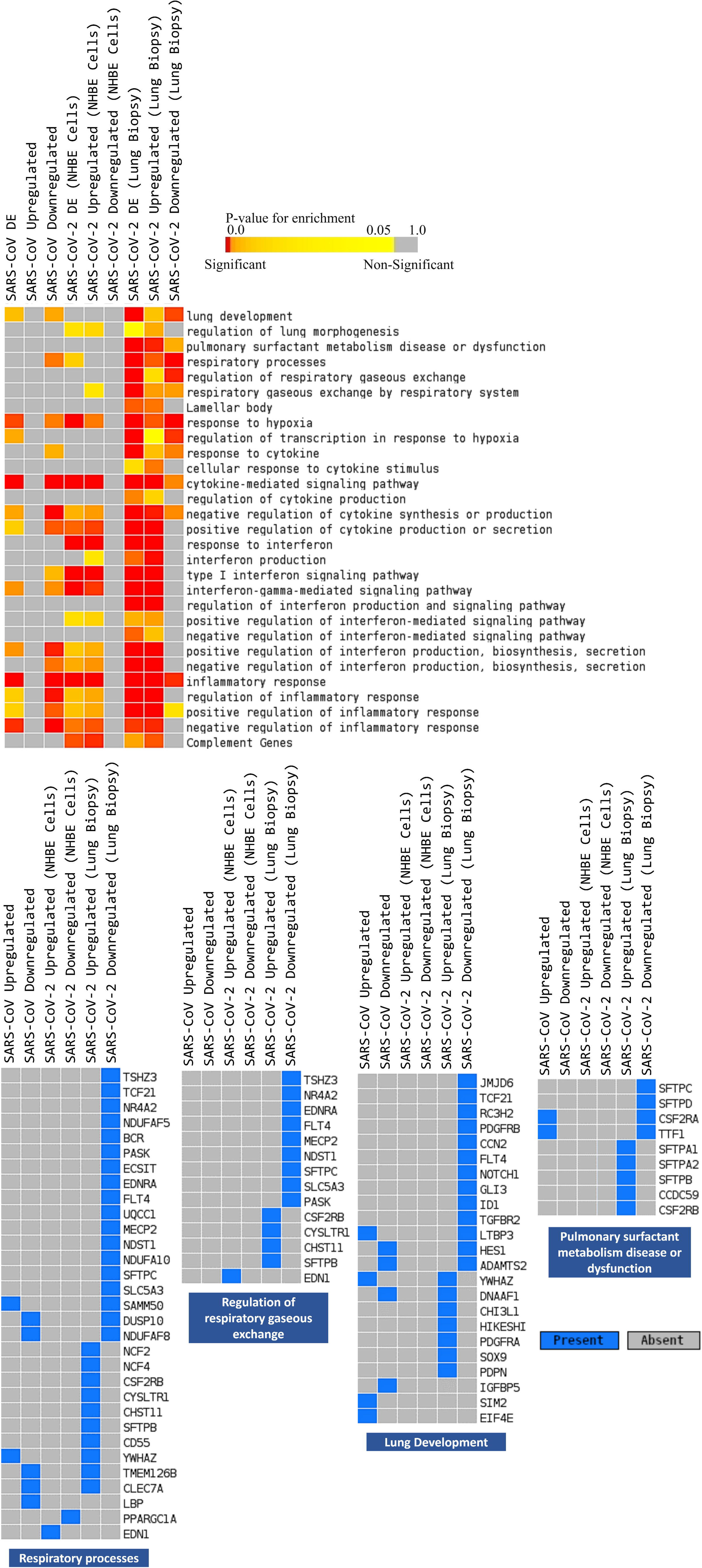
Enrichment analysis and comparison between deregulated genes and the genes of some selected processes in SARS-CoV, SARS-CoV-2 (NHBE cells) and SARS-CoV-2 (Lung biopsy) infections using combined module. Selected significant terms are represented in heatmap in upper panel. Color schemes are similar as Figure 1. All enriched terms and detail statistics are in Supplementary file 6. Lower panel heatmaps presents enriched genes for some selected terms from upper panel enrichment analysis. For individual processes, blue means presence (significantly differentially expressed gene) while grey means absence (not significantly differentially expressed genes for this module for this experimental condition).

### Genes in lung surfactant metabolism pathway are deregulated in COVID-19 patient’s lung

Pulmonary surfactant proteins play important role in maintaining the surface tension at the air-liquid interface in the alveoli for efficient gaseous exchange, and can also modulate immunological functions of lung’s innate immune cells to eliminate pathogens [53]. Moreover, their role in viral infections as well as in impeding inflammatory responses and clearance of apoptotic cells in lungs are previously reported [53]. Our above analysis as indicated that surfactant metabolism could be a target of this virus, we sought to check the routes of deregulations of this pathway in SARS-CoV-2 infected lung and how they are affecting the downstream signaling processes to impede normal lung function through impaired surfactant metabolism. To achieve this goal, we have mapped the differentially expressed genes in this pathway using Reactome pathway browser [38]. We looked deeply into the mechanisms of this pathway to elucidate the probable alterations happening in COVID-19 affected lung.

In normal lung, TTF1-CCDC59 complex can transactivate *SFTPB* and *SFTPC* gene expression which in turn play an important role by regulating the alveolar surface tension [54]. But in COVID-19 affected lung, *TTF1* and *SFTPC* genes are found to be downregulated, whereas *SFTPB* is upregulated (Figure 3). GATA6 transcription factor promotes the transcription of *SFTPA* gene [55] which is involved in immune and inflammatory responses along with lowering the surface tension in the alveoli [56] and both of them are found to be upregulated in SARS-CoV-2 infected lung while the GATA6 antagonist LMCD1 was downregulated (Figure 3). While the complex of CSF2RA and CSF2RB can bind GM-CSF to induce activation of macrophages [57] and degradation of STFPs in the alveolar macrophages [58]. We found *CFSF2RA* downregulated and *CSF2RB* upregulated in the lung of the COVID-19 affected patient (Figure 3). Pro-SFTPB and Pro-SFTPC are cleaved NAPSA, CTSH, PGA3-5 for producing active SFTPB and SFTPC [59, 60] but in the COVID-19 affected lung NAPSA was found deregulated (Figure 3) while the rest other enzymes were not found to be significantly differentially expressed (data not shown). Overall, the transcription of the surfactant genes, production of active surfactant proteins, and their turnover might be deregulated in the lung of the COVID-19 patient which could have result in the severe disease complicacies leading to their death. As these mechanisms are found to be aberrantly modulated upon SARS-CoV-2 infections, virus might be positively facilitating these anomalies.

**Figure 3:**
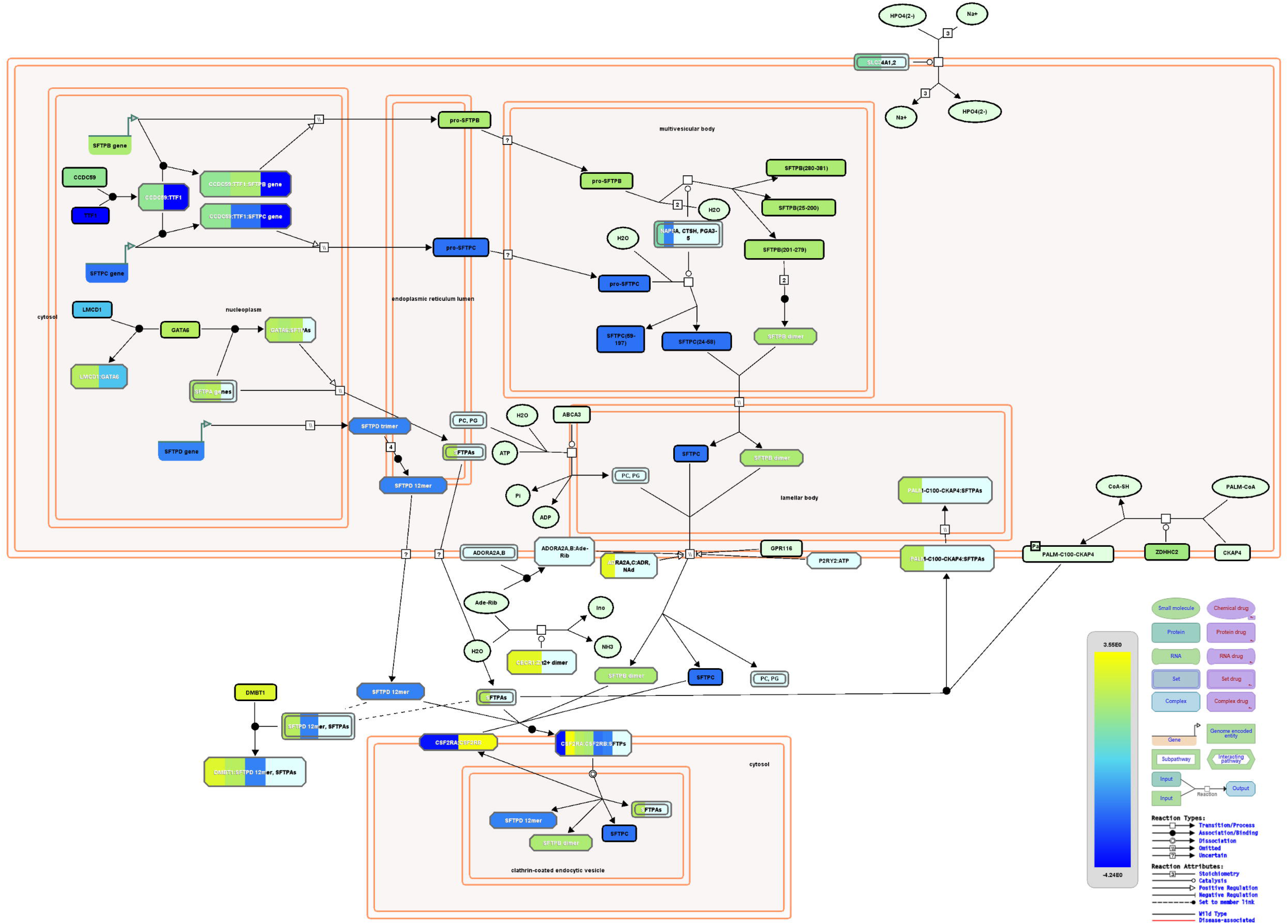
Schematic representation of lung surfactant metabolism pathway from Reactome pathway database. Color towards yellow indicates upregulation and while blue indicates downregulation.

### Host proteins those interact with virus are involved in different respiratory functions related pathways and diseases

In previously reported human coronavirus infections, SARS and MERS coronaviruses are often found to takeover host machineries, suppressing host immune responses and other important biological processes for their continued existence inside the infected cells [61]. We have performed functional enrichment analyses using the previously reported host proteins which can interact with SARS-CoV and SARS-CoV-2 proteins [33–35] to have better understanding in which pathways these proteins are involved and if there any connection with lung degenerative processes – a common complication that we observe in corona infected patient.

Upon the enrichment analyses, as expected, we observed several immune signaling pathways like-interleukin signaling, interferon signaling, apoptosis, inflammasomes etc. for both SARS-CoV and SARS-CoV-2 proteins (Figure 4A, 4C); however, this approach also revealed some vital pathways related to respiratory functions like- HIF-1 signaling, hypoxic and oxygen homeostasis regulation of HIF-1 alpha etc (Figure 4A, 4C). Several respiratory complications related pathways like- lung diseases, asthma, hyperoxia, respiratory failure, pulmonary hypertension etc. were found to be enriched from DisGenNet database module [41] (Figure 4B). All these enlighten that SARS-CoV-2 might be utilizing its proteins in modulating the host lung’s normal physiological and immune responses which can be further explored by linking these viral-host protein-protein interactions (PPI) in our previously identified essential lung processes.

**Figure 4:**
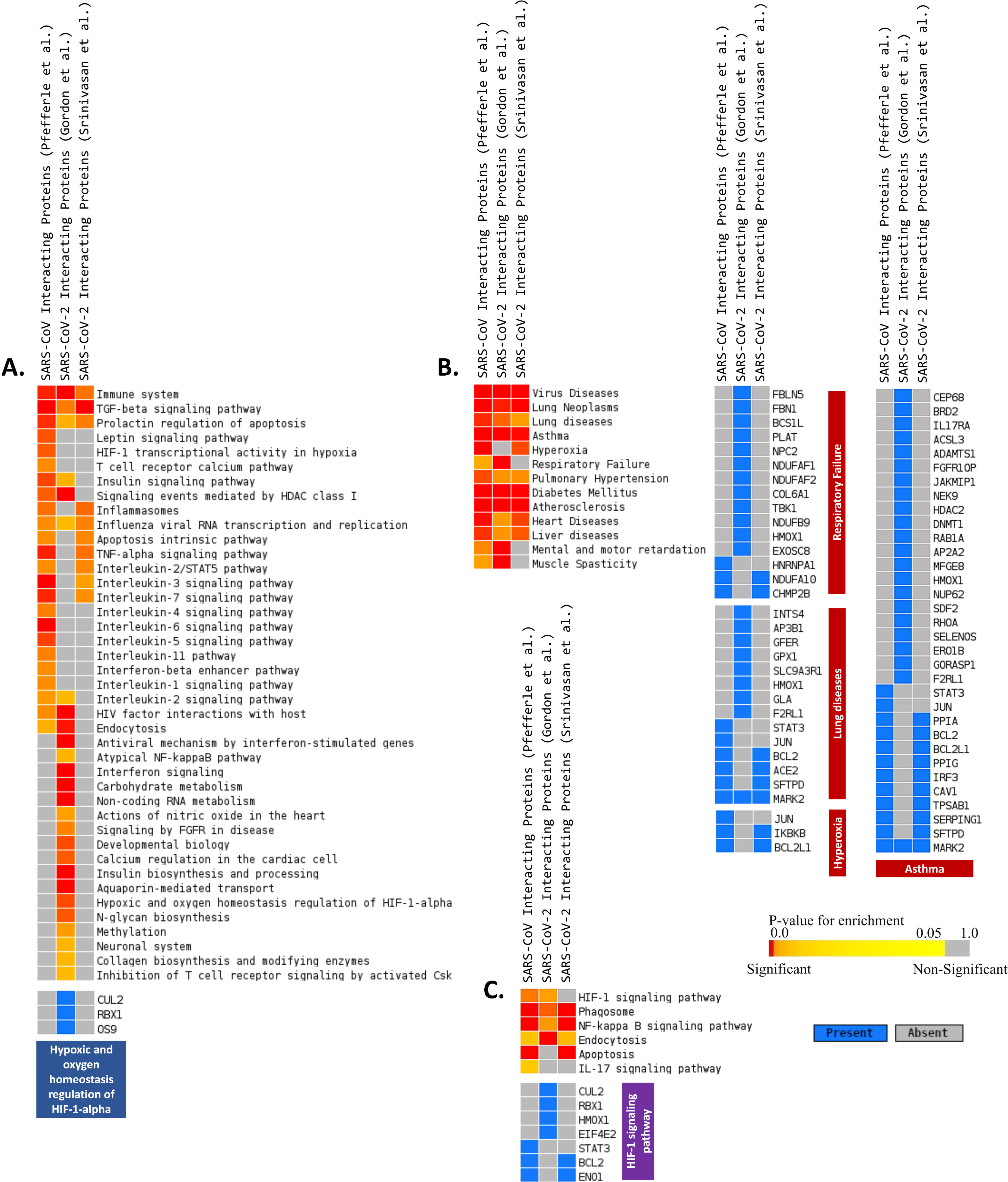
Enrichment analysis and comparison between host proteins interacting SARS-CoV proteins (Pfefferle et al.), SARS-CoV-2 proteins (Srinivasan et al.) and SARS-CoV-2 proteins (Gordon et al.) and the genes of some selected processes using **A.** Bioplanet pathway module, **B.** DisGeNet module, **C.** KEGG pathway module. Selected significant terms are represented in heatmap. All enriched terms and detail statistics are in Supplementary file 6. Color schemes are similar as Figure 1 and Figure 2.

### SARS-CoV-2 proteins and host epigenetic regulators can modulate the functionality of lung and other respiratory processes

As we have discovered that both deregulated genes in COVID-19 affected lung and SARS-CoV-2 interacting proteins are involved in different important respiratory functionalities, we then produced several functional networks with these deregulated genes, viral protein-host protein interactions, and host’s epigenetic regulators involved in those processes to shed insights on viral infection mediated deregulations and their resultant pathophysiological effects in COVID-19. We have mainly targeted four broad biological processes which can significantly affect the COVID-19 patients upon any dysregulation, namely-response to hypoxia, lung development, respiratory processes, and surfactant metabolism.

Numerous genes in the hypoxic responses and HIF-1 alpha signaling were found to be abruptly regulated in the SARS-CoV-2 infected lung (Supplementary Figure 5). PLAT, a tissue plasminogen activator, have profound roles in maintaining the normal homeostasis of lung and its aberrant regulation can lead to many lung injuries [62]. In the response to hypoxia process, in PPI map of differentially expressed genes of combined GOBP module ‘response to hypoxia’ and SARS-CoV-2 target genes [33], this PLAT protein was found to be directly targeted by viral protein ORF8 (Figure 5). Moreover, several indirect responses from the viral protein interactions were also revealed (Figure 5). SARS-CoV-2 M protein can target STOM which interacts with *SLC2A1*. *SLC2A1* can also be targeted by host miRNA miR-320a (Figure 5). ORF8 which interacts with OS9 can modulate EGLN1 and EGLN2. KCNMA1 is found to be interacting with host proteins ATP1B1 and PRKACA which interact with viral M and NSP13 proteins, respectively (Figure 5). ORF9c can modulate EDNRA indirectly through F2RL1 (Figure 5). PML functions might be altered through viral N-host MOV10 proteins’ interaction. Functions of SLCBA1 can be modulated by NSP13 through PRKACA (Figure 5). NSP7 can regulate ALDH3A1 which can affect CYP1A1 and NR4A2 (Figure 5). NSP5 interacting HDAC2 can curb PML and REST functionalities (Figure 5). NSP12 can modulate the a wide range of hypoxia functions related proteins like-TGFBR2, MECP2, MTHFR, CBFA2T3, EGR1, ANGPTL4 by interacting with the transcription factor TCF12 (Figure 5). NSP12 can deregulate the functions of CFLAR through RIPK1 interactions (Figure 5). Apart from these several host miRNAs can possibly also downregulate the expression of some genes, namely-miR-320a, miR-3188, miR-3661, miR-217, miR-421 and miR-429 (Figure 5). These viral mediated deviations found in the hypoxia responses might be a decisive factor behind the lung injury found in COVID-19 patients [63].

**Figure 5:**
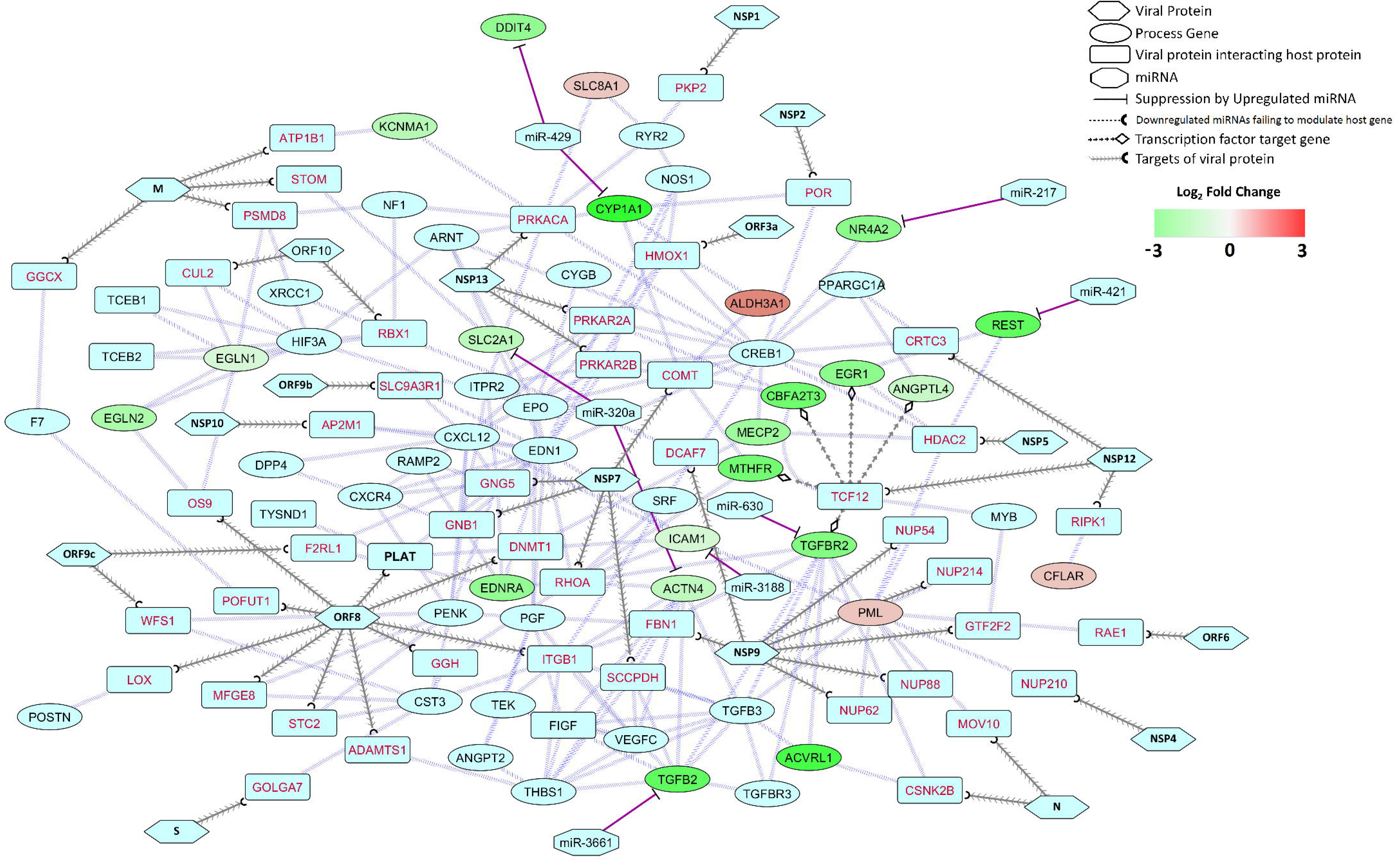
Network representing the interactions between genes in response to hypoxia process (combined module genes) along with SARS-CoV-2 proteins (Gordon et al.), and host miRNAs. Hexagon, ellipse, rounded rectangle, octagon represents viral proteins, process related genes, proteins that interacts viral proteins and host miRNAs, respectively. Blunted arrow indicates suppression by miRNAs, dotted arrow pointed with open half-circle indicates downregulated miRNAs failing to modulate host gene, arrowed line pointed with open half-circle indicates targets of viral proteins, and arrowed line pointed with open diamond indicates transcription factors of a gene.

In the lung development network, viral protein NSP13 are found to indirectly target RC3H2, SOX9, GLI3 through CEP350, PRKACA, PRKAR2A proteins (Figure 6). While GLI3 can also be modulated through several viral-host interactions like- ORF10-RBX1, NSP5-HDAC2, M-PSMD8 interaction axis (Figure 6). ORF8 can indirectly modulate ADAMTS2, CHI3L1, NOTCH1, TCF12, FLT4 etc (Figure 6). Transcription factor TCF12 is targeted by viral NSP12 protein which can in turn affect transcriptions of *PDGFRA*, *PDGFRB*, *TGFBR2*, *ID1*, *HES1*, *LTBP3* genes (Figure 6). NSP5 can modulate NOTCH1 through HDAC2 (Figure 6). Moreover, host miRNAs like- miR630 can downregulate TGFBR2; while miR-206, miR320a and miR-375 are downregulated so that SOX9 and WYHAZ are overexpressing (Figure 6). Virus could hamper several growth factor signaling which are crucial for various lung injury repair mechanisms [64, 65].

**Figure 6:**
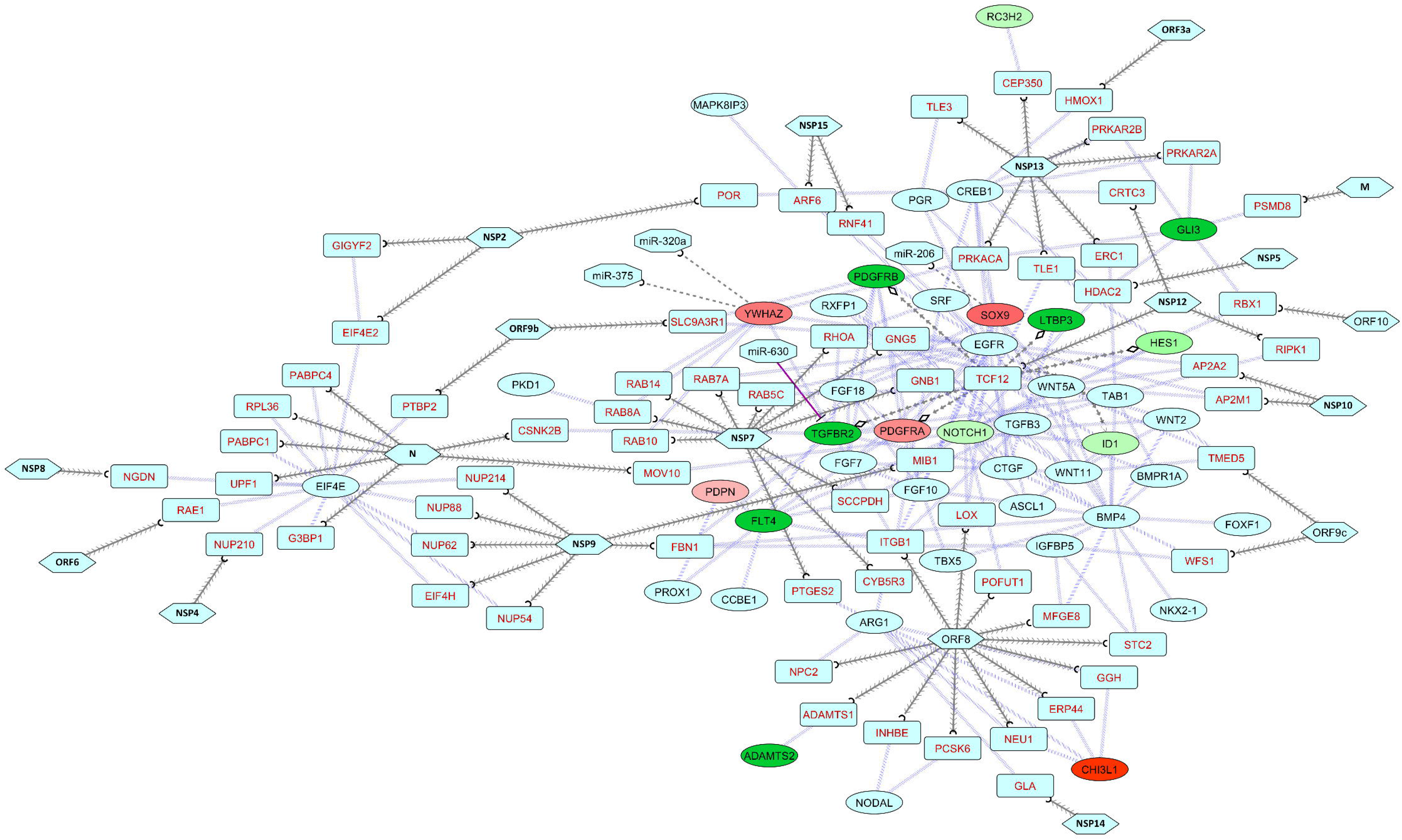
Network representing the interactions between genes in lung development process (combined module genes) along with SARS-CoV-2 proteins (Gordon et al.), and host miRNAs. Legends are similar as Figure 5.

From the respiratory process network we can delineate that several transcription factors are deregulated which are associated in transcribing ECSIT that is directly targeted by viral ORF9c and indirectly by viral ORF8, NSP7 (Figure 7). Moreover, ECSIT itself is directly targeted by ORF9c (Figure 7). NSP12 can modulate SFTPB, SFTPC, SLC5A3, DUSP10, SAMM50 etc. by targeting TCF12 (Figure 7). Also, in this network we have observed suppressive actions of miRNAs- miR-206, miR-217, miR-375 on NR4A2 and NDST1; as well as upregulation of YWHAZ due to probable downregulation of miR-320a (Figure 7). Virus might be dampening the host immune response in lung by targeting ECSIT [66], as well as could dull the respiratory gaseous exchange in lung by negatively modulating the surfactant proteins [67].

**Figure 7:**
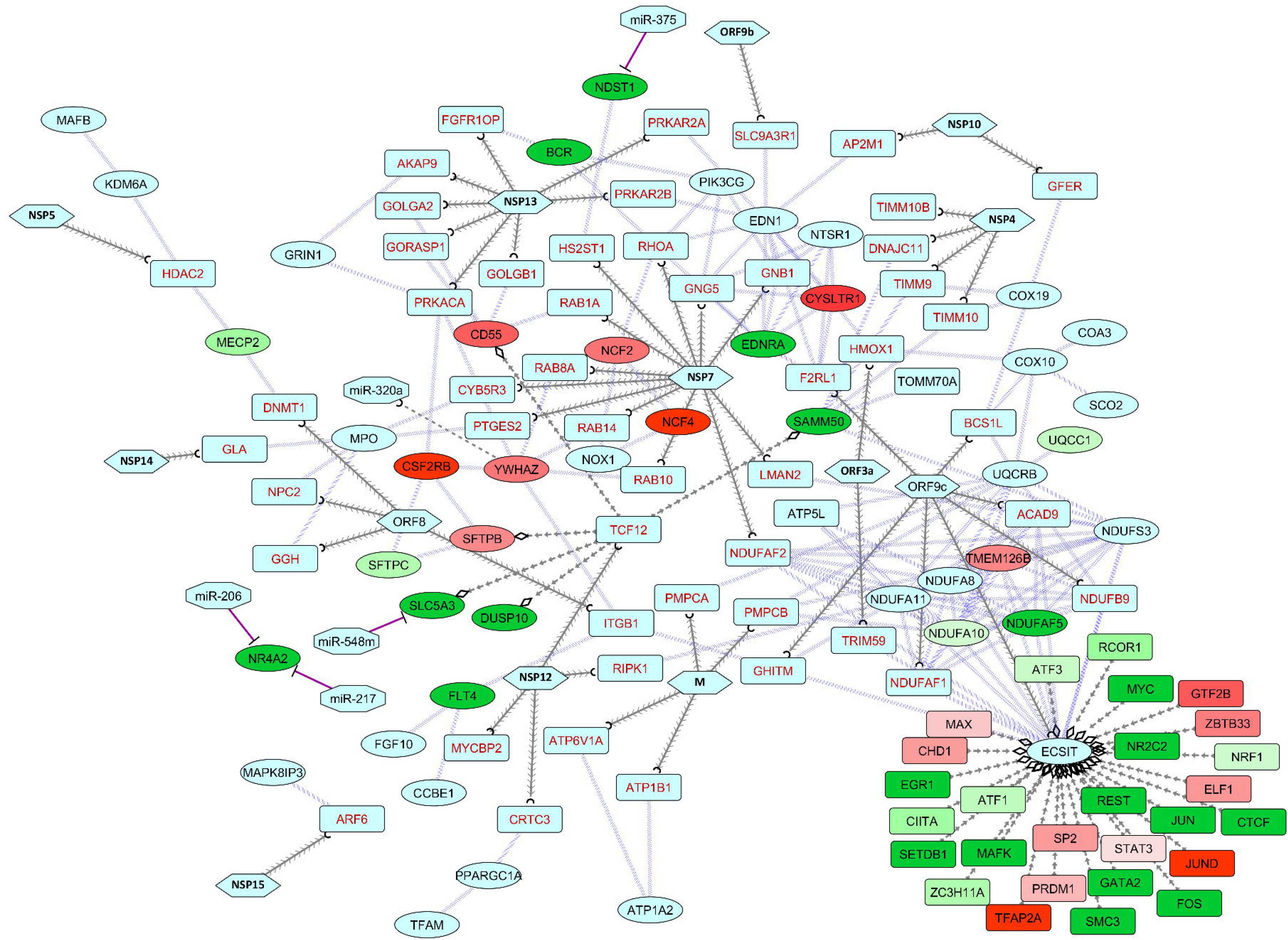
Network representing the interactions between genes in respiratory processes (combined module genes) along with SARS-CoV-2 proteins (Gordon et al.), and host miRNAs. Legends are similar as Figure 5.

Surfactant metabolism is found to be modulated by not only viral proteins but also through the aberrant host response (Figure 8). Viral proteins NSP12 and NSP5 can target transcription factor TCF12 and epigenetic regulator HDAC2, respectively which in turn are modulating important members of surfactant metabolism process like- TTF1, CCDC59, SFTPB, SFTPC, CSF2RA, CSF2RB, NAPSA, SFTPD, DMBT1 etc (Figure 8). These can also be modulated through CKAP4 which are targeted by viral M, NSP2, NSP9, E, ORF8 proteins (Figure 8). Furthermore, we have observed that viral M and S proteins can interact with the proteins of this process both directly and indirectly (Supplementary figure 6). Host miRNAs miR-421 are found to be downregulating LMCD1; while miR-137, miR-375, miR-429 fail to modulate CCDC59 and GATA6 because of their probable inactivation/suppression by differential expression of its regulating TFs (Figure 8). As *SFTPD* and *SFTPC* are downregulated along with several regulatory partners, their primary function of immunomodulation and efficient air exchange in lung [68, 69] might be seriously hindered by the viral proteins; which could further lead to abnormal pathogenic lung injury [70].

**Figure 8:**
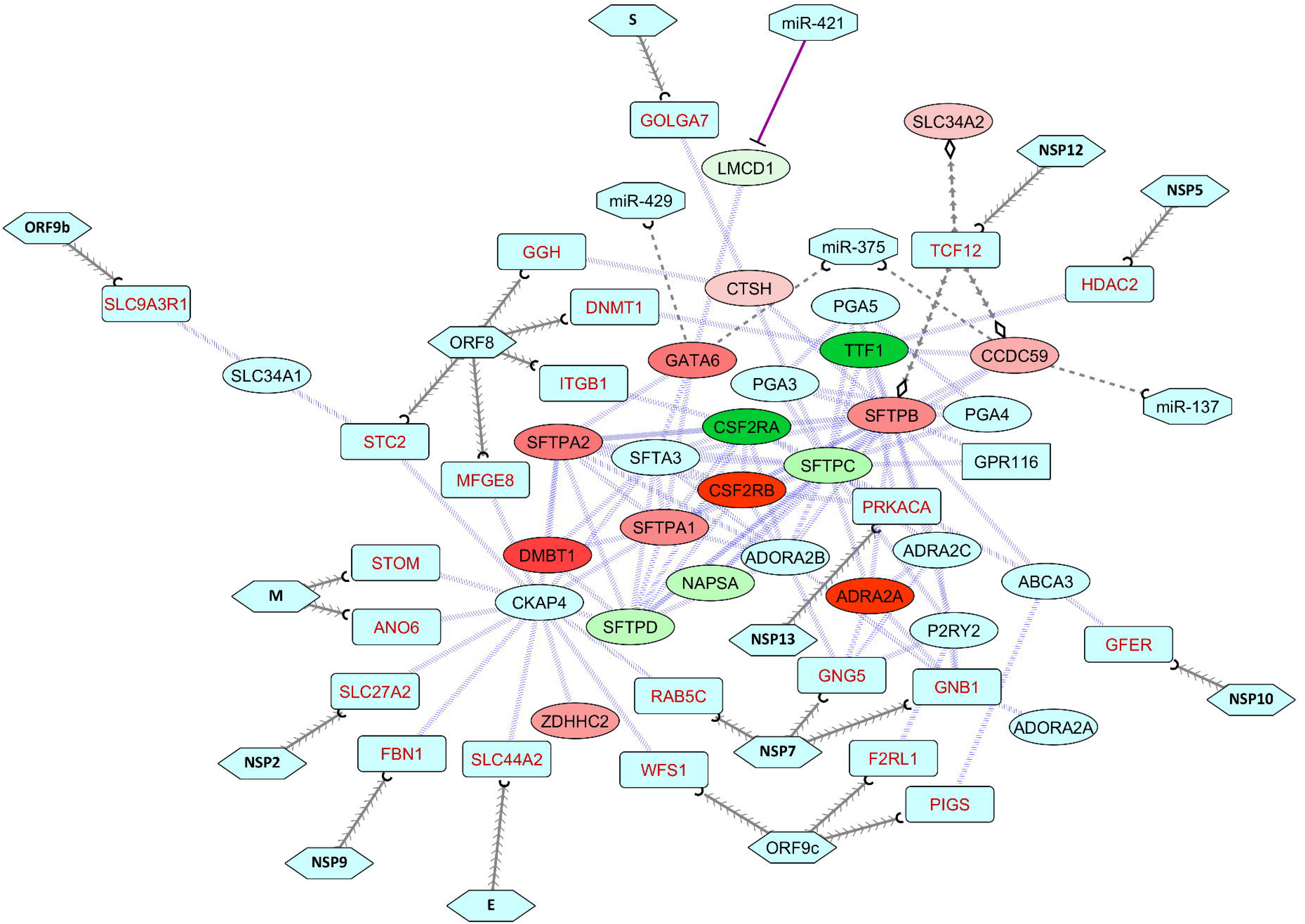
Network representing the interactions between genes in surfactant metabolism along with SARS-CoV-2 proteins (Gordon et al.), and host miRNAs. Legends are similar as Figure 5.

Several epigenetic factors like- HDAC2, DNMT1, CUL2, MOV10, RBX1 and TLE1 which are involved in these processes can be targeted by different viral proteins, through which these abovementioned processed can be significantly altered. Epigenetic modulator are key modulators in balancing the normal lung pathobiology and anomalies of these regulations can lead to many lung diseases [71]. HDAC2 and DNMT1 have significant roles in chronic obstructive pulmonary disease (COPD) progression [72, 73].

## Discussion

Though there are a wide variety of symptoms and clinical features seen in the COVID-19 patients, almost every mild to critically affected patients showed respiratory and breathing complications which ranges from pneumonia to acute lung injury [74]. Severely ill patients are mostly supported by artificial ventilation as no therapeutic drugs are discovered yet for mitigating these complications due to the missing understandings on the molecular aspects of lung related abnormalities in COVID-19. We have identified some important respiratory and lung related processes in which we got several genes are deregulated upon the progression of SARS-CoV-2 infection. So, we prioritized these pathways to check how viral proteins might be playing a role in these aberrant regulations and searched for potential therapies/drugs for the treatment to lessen the resultant effects of the aberration.

We have compared the deregulated genes from SARS-CoV-2 infected lung with the differentially expressed genes from SARS-CoV-2 infected cell line and SARS-CoV infection, to see how host lung and SARS-CoV-2 are responding upon the infections. We found that a significant variations in the number of total deregulated genes between these. Other report also suggests the deregulation of a huge number of genes when expression anylyses are performed using host’s infected cells [75]. That means cell lines infected with SARS-CoV-2 might not provide the complete outcomes of the infections. We also checked and verified the quality of our normalized reads to nullify the risk of probable poor sequencing and abnormal read normalization results; all these recommend the suitability of our generated differential expression analysis.

Several SARS-CoV-2 entry receptor (ACE2)/entry associated proteins (TMPRSS2, BSG, CTSL, DPP4) were previously discussed [76]; all of which are readily expressed in the lung, except ACE2 (Supplementary Figure 7A). While analyzing the expression profiling in SARS-CoV-2 infected lung, we discovered a quite unexpected scenario. Astoundingly, in COVID-19 affected lung, *ACE2* was upregulated while the genes of the entry associated proteins were downregulated (Supplementary Figure 7B). More COVID-19 patient data have to be analyzed in concluding this striking feature.

“Cytokine storm” is a much discussed phenomenon in previously reported pathogenic human coronavirus infections which can be lethal for the host for the destruction of its own respiratory systems [77]. Similar responses can also occur in SARS-CoV-2 infection [78]. We have also observed that several inflammatory and antiviral responses are readily deregulated in the SARS-CoV-2 infected lung which could lead to the abnormalities in overall respiratory functions.

Apart from this, our study also identified dysregulations in several non-cytokine mediated lung function related vital processes, namely- response to hypoxia, lung development, respiratory process, and surfactant metabolism; aberrations in these pathways leading to lung pathobiology in COVID-19. Networks generated combining the viral-host protein interactions information suggest that viral proteins might be actively involved in these deregulations along with several host factors which were also decontrolled due to the viral infections.

Hypoxic conditions are common in respiratory infections due to reduced inhalation of oxygen [79]. Several genes which are involved in hypoxia induced responses are found to be deregulated (Figure 5) in COVID-19 affected lung. ALDH3A1 can protect airway epithelial cells from destruction [80]. Though it is upregulated its functionalities can be impeded through viral proteins (Figure 5). CFLAR functions in shutting down apoptotic responses by interacting with RIPK1 [81] but viral interactions with RIPK1 might prevent it (Figure 5). Virus can induce the transcription of ANGPTL4 by utilizing TCF12 transcription factor (Figure 5) which in turn could cause pulmonary tissue damage [82]. Viral protein could promote the activity of EGLN1 and EGLN2 (Figure 5) to suppress the transcription of HIF-induced genes in hypoxia [83]; on the other hand the regulation through these proteins might be hampered due to viral ineractions and the constant overexpression of HIF mediated inflammatory genes could also occur, which could lead to inflammation induced lung damage. Severe hypoxia induced responses can occur through the downregulation of MECP2 [84] in SARS-CoV-2 patients (Figure 5). In hypoxia, REST is induced and can act as a negative regulator of gene expression to maintain a balance between different processes [85] which is here found downregulated and can be targeted through host miRNA miR-421 in COVID-19 (Figure 5). GLUT1 (*SLC2A1* gene) promotes increased glucose transport into hypoxic cells for its prolonged adaptation during this condition [86], but it is found to be downregulated in lung of COVID-19 patients and this can occur through miR-320a (Figure 5). During hypoxia, TGF-beta signaling regulates inflammation and vascular responses [87]; but attenuated TGFβ expression might lead to disease severity (Figure 5). EGR1 transcription factor can be modulated through viral proteins (Figure 5) in preventing hypoxia induced EGR1 activation of HIF-1 alpha [88], which could curb the whole hypoxia induced survival responses. Moreover, defective hypoxia response can occur due to the overactivity of EGR1 that can result in anomalous thrombosis [89]. Also, PLAT which is a crucial factor in splitting down clots [90], this function might be directly altered by SARS-CoV-2 ORF8. Many COVID-19 patients are reported to have pulmonary embolism and thrombosis [91], this hypoxic responses could suggest the probable routes of this.

Similarly, several lung development and respiratory process genes/proteins were also found to be deregulated in the COVID-19 patient’s lung (Figure 6, 7). SOX9 which is considered an important regulator for the recovery from acute lung injury [92], can be down-modulated by the virus (Figure 6). Though the host miRNAs cannot target the YWHAZ which is a pro-survival protein [93] and also can induce expression surfactant protein A2 [94], it can be indirectly modulated through viral protein NSP7 (Figure 6). Likewise, viral protein ORF8 can indirectly modulate CHI3L1 (Figure 6) which can suppress lung epithelium injury [95]. Virus can block TCF12 and inhibit the expression of PDGFRA and PDGFRB (Figure 6) which could play a role in lung maturation and injury response [96]. GLI3, which exerts essential role in developing the lung, and regulating the innate immune cells [97], is downregulated in COVID-19 lungs (Figure 6). LTBP3 can promote lung alveolarization [98] but can be modulated by SARS-CoV-2 protein NSP12 (Figure 6). Aberration in NOTCH signaling contributes significantly in various lung diseases [99], and in SARS-CoV-2 infection NOTCH1, HES1 is found to be downregulated (Figure 6). CD55, a member of the complement system which plays crucial role in host defense in airway epithelium [100], can be modulated by viral protein NSP12 (Figure 7). CYSLTR1 is upregulated in SARS-CoV-2 infection (Figure 7), which is correlated to COPD [101]. This protein can be modulated by viral ORF9c and ORF3a (Figure 7). ECSIT found to be directly targeted by viral ORF9c protein (Figure 7) to stop the ECSIT mediated antiviral innate immune response [66]. Several mitochondrial genes like- *NDUFA10*, *NDUFAF5*, *SAMM50* are found downregulated in COVID-19 affected lung (Figure 7); as mitochondria plays important role in cellular respiration and lung diseases [102], aberration of these might also lead to lung related complicacies. DUSP10 can regulate aberrant inflammatory response upon viral infections [103], but in SARS-CoV-2 infected lung this gene is downregulated which might have occurred through the interaction of NSP12-TCF12 interactions (Figure 7).

Pulmonary surfactant proteins are lipoproteins which mainly functions to lower the alveolar surface tension [104] as well as they can elicit some immune stimulatory roles against some respiratory pathogens [105]. Among the surfactant proteins, SP-A and SP-D mainly evoke immune responses while SP-B and SP-D play roles in maintaining the efficient respiratory gaseous exchanges [69]. Several lung diseases like- asthma, acute respiratory distress syndrome (ARDS), COPD etc. are reported to be associated with aberration in the functionalities of surfactant proteins [70, 106]. In SARS-CoV-2 affected lung, production of the surfactant proteins are found to be deregulated (Figure 8). Viral protein NSP5 can recruit HDAC2 and downregulate the expression of *TTF1* which is needed for the expression of SP-B and SP-C (Figure 8). SP-A and SP-D can be targeted indirectly by several viral proteins (Figure 8) which can lead to the aberrant production of surfactant proteins in lungs, thus might complicating the disease condition. CSF2RA and CSF2RB complex modulate the surfactant recycling, thus maintains the overall balance of surfactant content [58]. We found *CSF2RA* is deregulated in SARS-CoV-2 infected lung which could lead in the aberration of surfactant recycling (Figure 8). While performing the enrichment analysis, we witnessed that several cholesterol biosynthesis pathways are significantly downregulated in COVID-19 patient’s lung cells (Figure 1E) which could lead to low accumulation of phospholipids in lungs. As phosphatidylcholine (PC) and phosphatidylglycerol (PG) are the principal phospholipids of surfactant proteins [107], impeded production of lipids will make surfactant proteins non-functional.

Considering all these, lung surfactants might be useful in the treatment of COVID-19 patients, as lung surfactant replacement therapies were previously reported to be successful in other respiratory infections and acute lung injury to reduce the lung damage of the patients [104, 108, 109] and also our analysis found it as an alternative from enrichment of related process deregulated genes (data not shown). Also, other drugs like- respiratory stimulants for COPD [110], sargramostim for treating pulmonary alveolar proteinosis [111], and oseltamivir in curing influenza-related lower respiratory tract complications [112] showed potential improvements in lung’s and respiratory system’s overall condition. Further *in vitro* and *in-vivo* research may be turn out to be useful for the patients who cannot tolerate painful ventilator.

From our results, we can suggest that surfactant proteins production along with other respiratory responses in lung could be deregulated. Along with the antiviral drugs to mitigate the viral responses, drugs which can improve the lung conditions in COVID-19 patients could also be considered as a treatment option for the patients. Our generated model can be useful for further experiments, as well as more insights will be gathered upon the use of surfactant therapy in laboratory using SARS-CoV-2 infected model organisms.

## Conflict of Interest

The authors declare that the research was conducted in the absence of any commercial or financial relationships that could be construed as a potential conflict of interest. The authors declare no conflict of interest.

## Author’s Contribution

ABMMKI conceived the project, designed the workflow and performed the analyses. Both authors interpreted the results and wrote the manuscript. Both authors read and approved the final manuscript.

## Funding

This project was not associated with any internal or external source of funding.

## Data Availability Statement

Analyses generated data are deposited as supplementary files.

## Supplementary Figure legends

**Supplementary Figure 1:** Venn diagrams for comparing the differences between **A.** Deregulated genes in SARS-CoV, SARS-CoV-2 (NHBE cells), and SARS-CoV-2 (lung biopsy) infections, **B.** upregulated genes in SARS-CoV, downregulated genes in SARS-CoV-2 (NHBE cells), and upregulated genes in SARS-CoV-2 (lung biopsy) infections, **C.** upregulated genes in SARS-CoV, upregulated genes in SARS-CoV-2 (NHBE cells), and downregulated genes in SARS-CoV-2 (lung biopsy) infections, **D.** upregulated genes in SARS-CoV, downregulated genes in SARS-CoV-2 (NHBE cells), and downregulated genes in SARS-CoV-2 (lung biopsy) infections, **E.** upregulated genes in SARS-CoV, upregulated genes in SARS-CoV-2 (NHBE cells), and upregulated genes in SARS-CoV-2 (lung biopsy) infections, **F.** downregulated genes in SARS-CoV, upregulated genes in SARS-CoV-2 (NHBE cells), and upregulated genes in SARS-CoV-2 (lung biopsy) infections, **G.** downregulated genes in SARS-CoV, downregulated genes in SARS-CoV-2 (NHBE cells), and upregulated genes in SARS-CoV-2 (lung biopsy) infections, **H.** downregulated genes in SARS-CoV, upregulated genes in SARS-CoV-2 (NHBE cells), and downregulated genes in SARS-CoV-2 (lung biopsy) infections, **I.** downregulated genes in SARS-CoV, SARS-CoV-2 (NHBE cells), and SARS-CoV-2 (lung biopsy) infections.

**Supplementary Figure 2:** Deregulated genes of selected terms from Figure 1 in SARS-CoV, SARS-CoV-2 (NHBE cells) and SARS-CoV-2 (Lung biopsy) infections. Genes of selected significant terms are represented here. For individual processes, blue means presence (differentially expressed gene of the module term) while grey means absence (not differentially expressed in the experimental condition in that module term). Processes in orange, green, purple, red, cyan color background represent Bioplanet, HumanCyc, GOBP, Reactome, DisGeNet enriched terms, respectively.

**Supplementary Figure 3:** Deregulated genes of additional terms from combined module based enrichment in SARS-CoV, SARS-CoV-2 (NHBE cells) and SARS-CoV-2 (Lung biopsy) infections. Genes of selected significant terms are represented here. For individual processes, blue means presence while grey means absence (color code as in Supplementary Figure 2).

**Supplementary Figure 4:** Expression profiles of genes in different lung associated processes obtained from enrichment analysis with combined module. Color towards red indicates more differential upregulation while color towards green indicates more differential downregulation; yellow color suggesting less expression change compared to control.

**Supplementary Figure 5:** Network representing the interactions between genes in surfactant metabolism along with SARS-CoV-2 proteins (Srinivasan et al.), and host miRNAs. Legends are similar as Figure 5.

**Supplementary Figure 7:** Expression profiles of SARS-CoV-2 receptor/associated proteins for entry into the target cell in **A.** Different organs of normal human, **B.** SARS-CoV-2, SARS-CoV infections. For FPKM scale, color towards blue means higher expression while color towards grey indicates lower level of expression. For Log_2_ fold change scale, color towards red indicates upregulation while color towards green indicates downregulation; yellow color suggesting no significant expression change.

## List of supplementary files

**Supplementary file 1:** Log_2_ Normalized expression data of control and SARS-CoV infection (GEO accession: GSE17400).

**Supplementary file 2:** Raw read counts expression of RNA-seq data (GEO accession: GSE147507) of control and SARS-CoV-2 infection.

**Supplementary file 3:** Quality analysis results of processed normalized RNA-seq data using “arrayQualityMetrics”.

**Supplementary file 4:** List of the combined modules along with the genes and their associated original term and sources.

**Supplementary file 5:** List of host proteins which interact with SARS-CoV, SARS-CoV-2 proteins.

## Notes

### Competing Interest Statement

The authors have declared no competing interest.

