## Supplementary figures and images for "Lung biopsy cells transcriptional landscape from COVID-19 patient stratified lung injury in SARS-CoV-2 infection through impaired pulmonary surfactant metabolism"

### Supplementary Figure 1

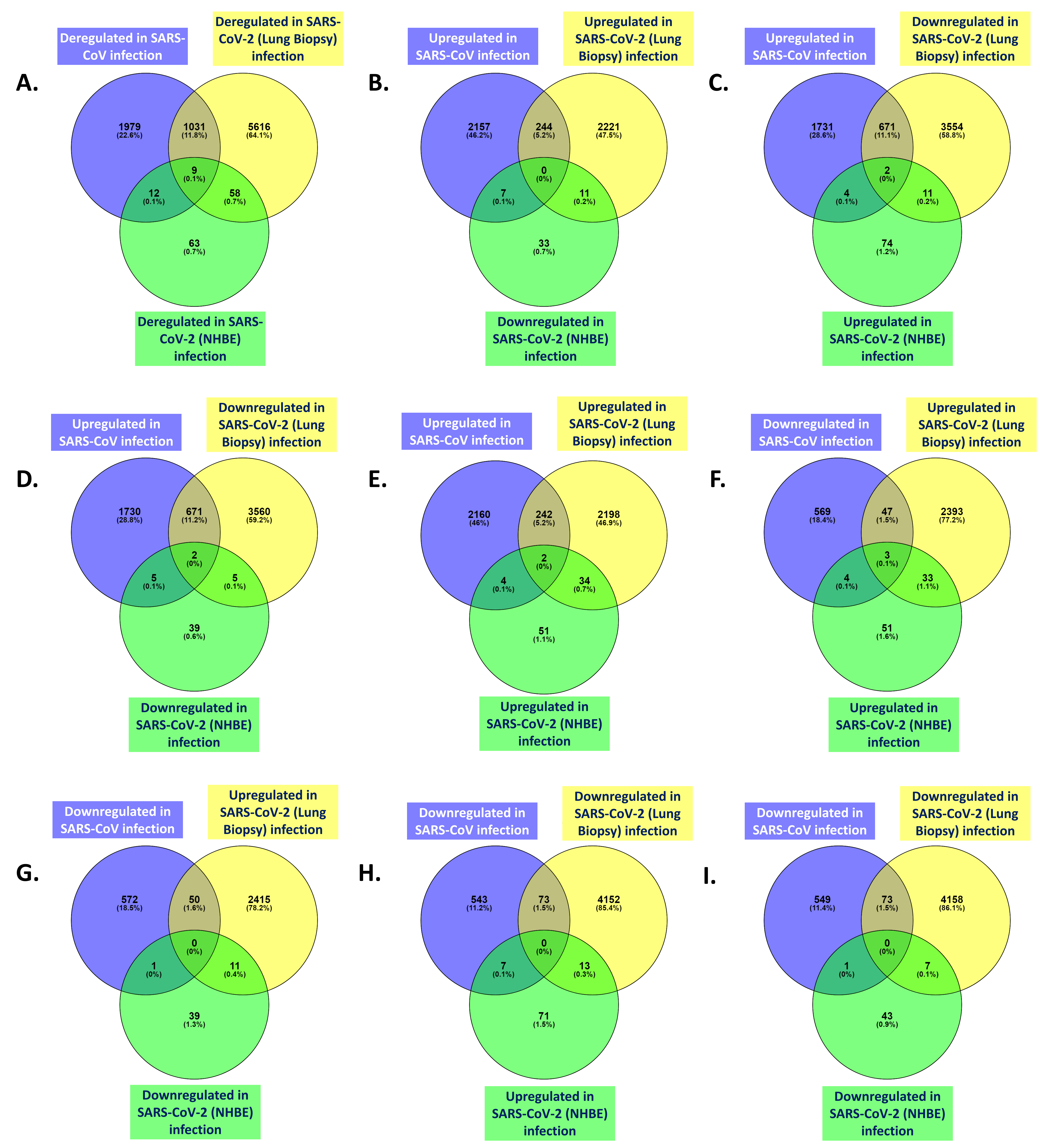

### Supplementary Figure 2

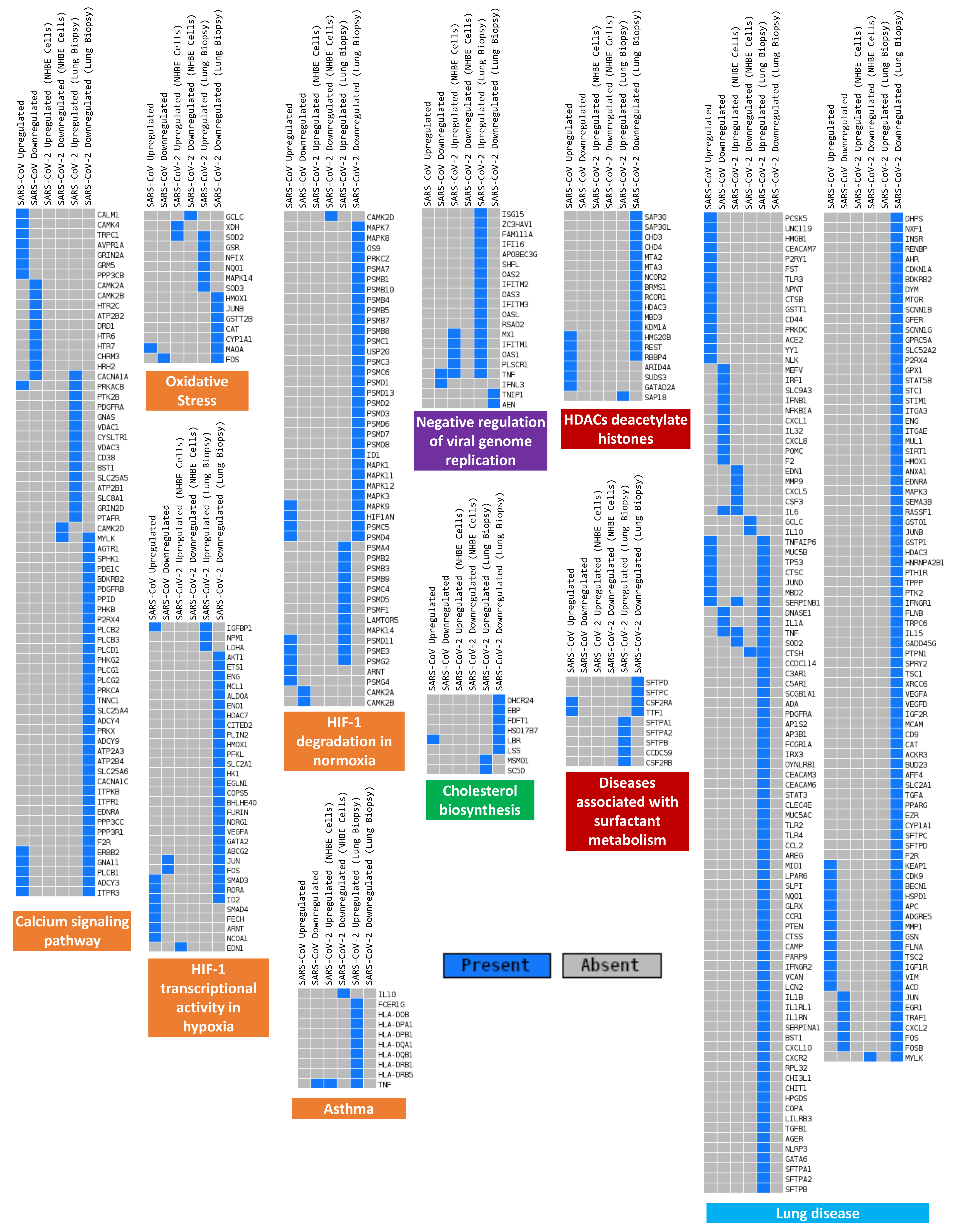

### Supplementary Figure 3

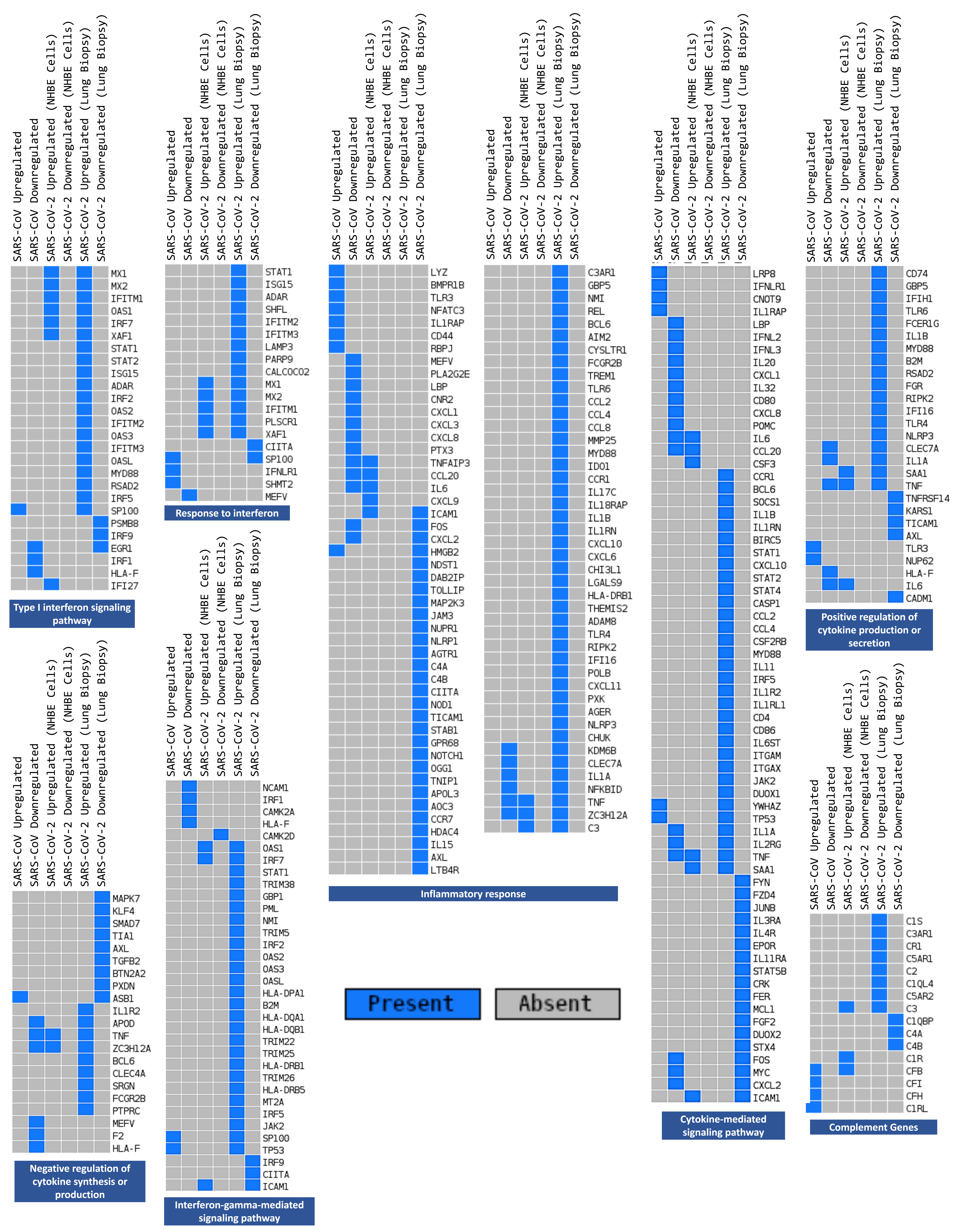

### Supplementary Figure 4

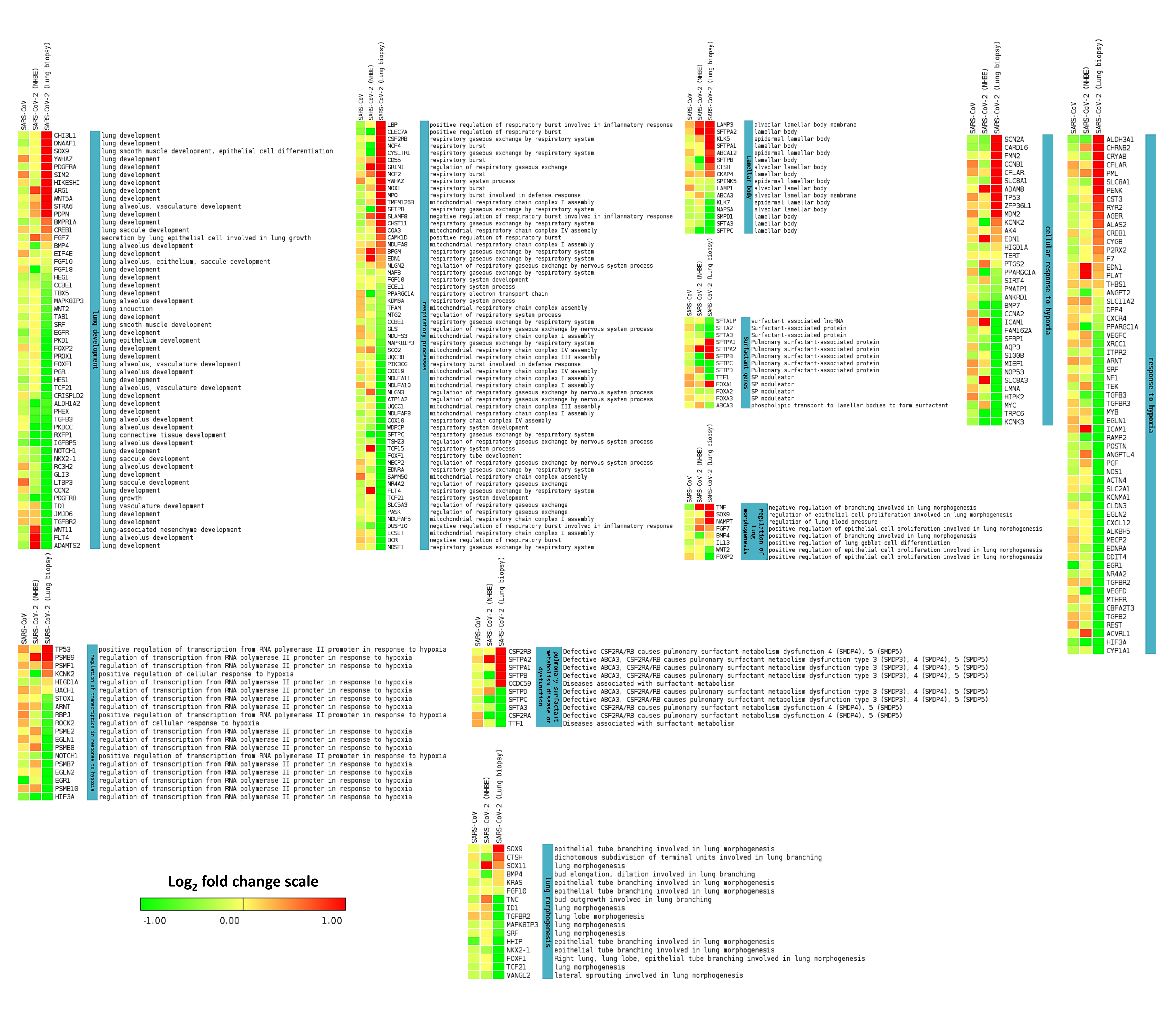

### Supplementary Figure 5

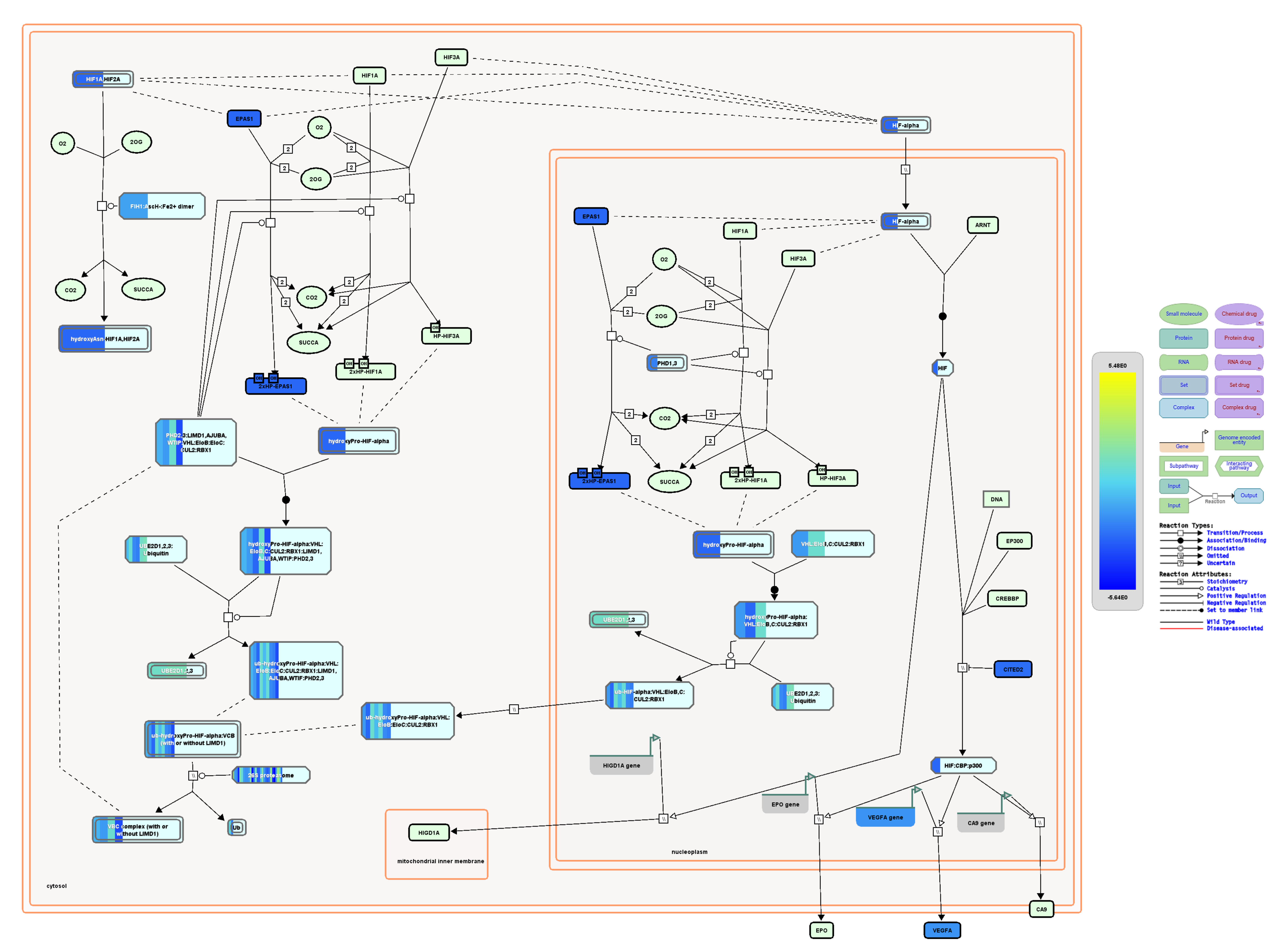

### Supplementary Figure 6

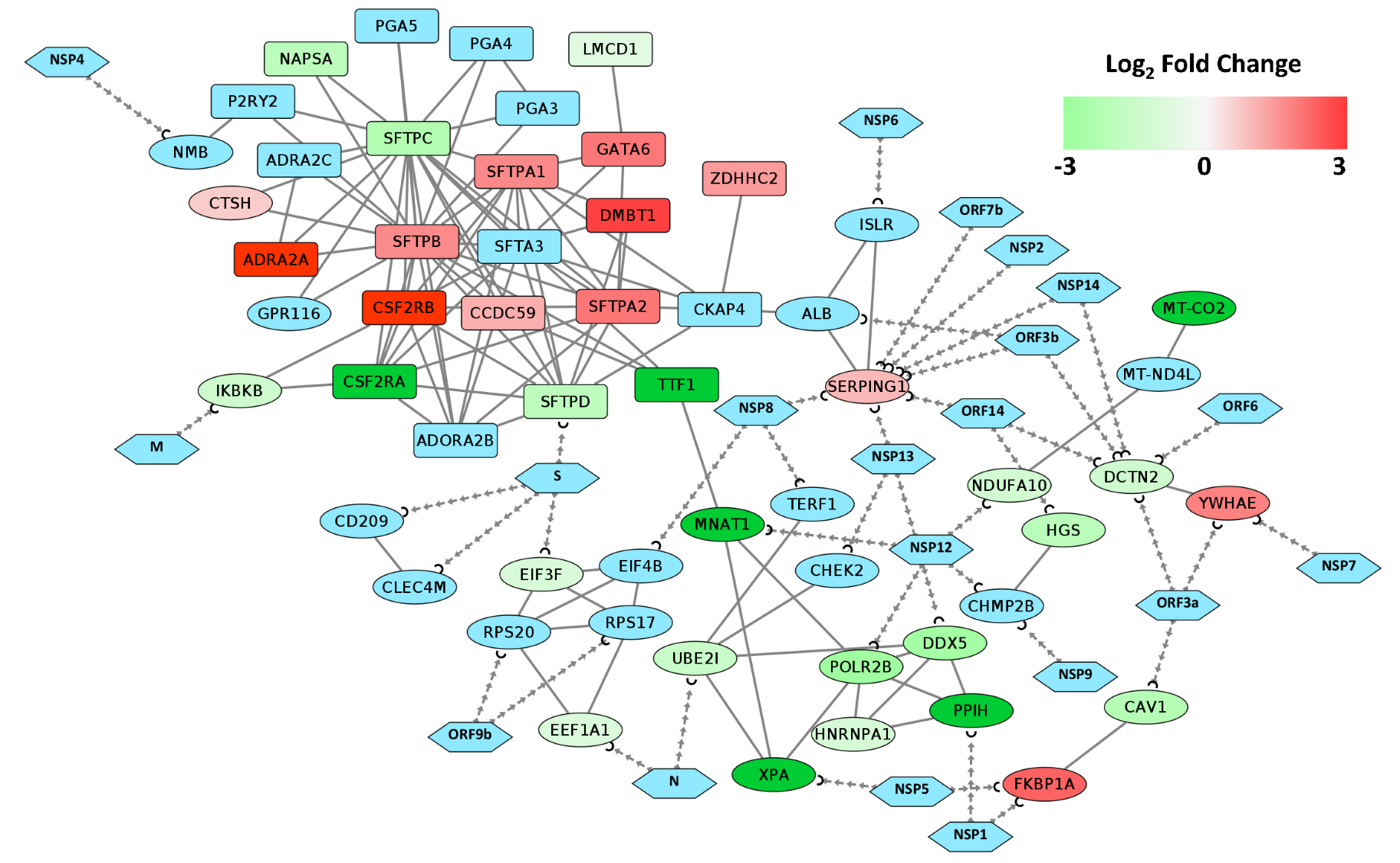

### Supplementary Figure 7

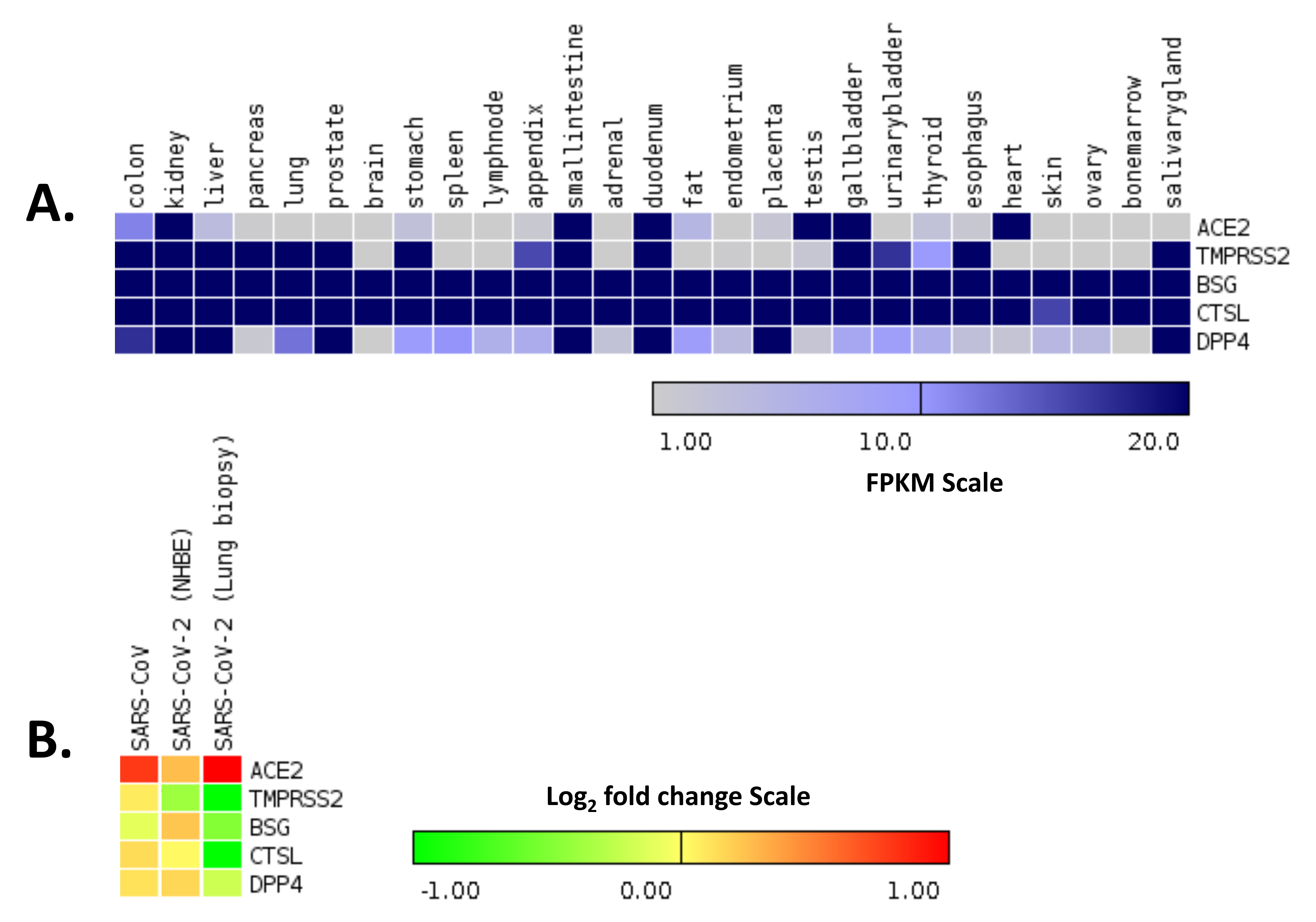
