## Supplementary File 3 for "Lung biopsy cells transcriptional landscape from COVID-19 patient stratified lung injury in SARS-CoV-2 infection through impaired pulmonary surfactant metabolism"

**Supplementary file 3: Quality analysis results of processed normalized RNA-seq data using “arrayQualityMetrics”.**

**Section 1: Between array comparison**

**[- Figure 1: Distances between arrays.](javascript:%20toggle('hm'))**

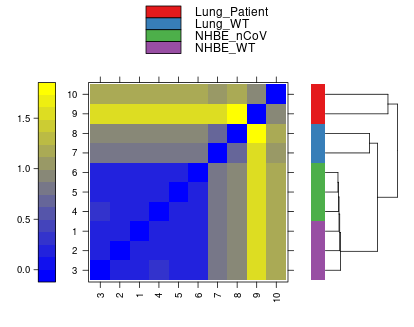

**Figure 1** shows a false color heatmap of the distances between arrays. The color scale is chosen to cover the range of distances encountered in the dataset. Patterns in this plot can indicate clustering of the arrays either because of intended biological or unintended experimental factors (batch effects). The distance *d_ab_* between two arrays *a* and *b* is computed as the mean absolute difference (L_1_-distance) between the data of the arrays (using the data from all probes without filtering). In formula, *d_ab_* = mean | *M_ai_ - M_bi_* |, where *M_ai_* is the value of the *i*-th probe on the *a*-th array. Outlier detection was performed by looking for arrays for which the sum of the distances to all other arrays, *S_a_* = Σ*_b_* *d_ab_* was exceptionally large. No such arrays were detected.

**[- Figure 2: Outlier detection for Distances between arrays.](javascript:%20toggle('out%20hm'))**

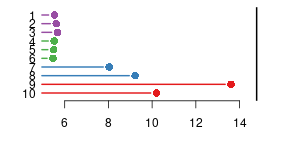

**Figure 2** shows a bar chart of the sum of distances to other arrays *S_a_*, the outlier detection criterion from the previous figure. The bars are shown in the original order of the arrays. Based on the distribution of the values across all arrays, a threshold of 14.8 was determined, which is indicated by the vertical line. None of the arrays exceeded the threshold and was considered an outlier.

**[- Figure 3: Principal Component Analysis.](javascript:%20toggle('pca'))**

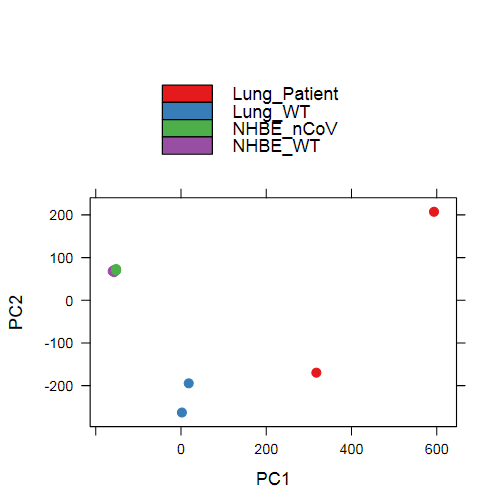

**Figure 3** shows a scatterplot of the arrays along the first two principal components. You can use this plot to explore if the arrays cluster, and whether this is according to an intended experimental factor, or according to unintended causes such as batch effects. Move the mouse over the points to see the sample names.

Principal component analysis is a dimension reduction and visualisation technique that is here used to project the multivariate data vector of each array into a two-dimensional plot, such that the spatial arrangement of the points in the plot reflects the overall data (dis)similarity between the arrays.

**Section 2: Array intensity distributions**

**[- Figure 4: Boxplots.](javascript:%20toggle('box'))**

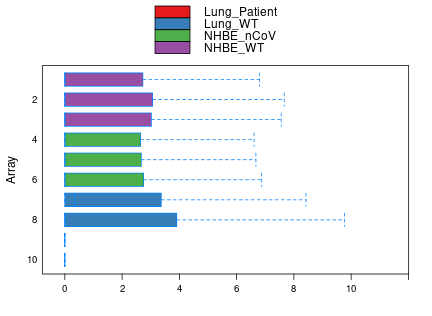

**Figure 4** shows boxplots representing summaries of the signal intensity distributions of the arrays. Each box corresponds to one array. Typically, one expects the boxes to have similar positions and widths. If the distribution of an array is very different from the others, this may indicate an experimental problem. Outlier detection was performed by computing the Kolmogorov-Smirnov statistic *K_a_* between each array's distribution and the distribution of the pooled data.

**[- Figure 5: Outlier detection for Boxplots.](javascript:%20toggle('out%20box'))**

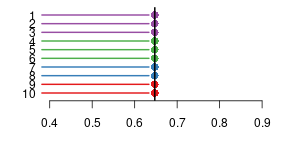

**Figure 5** shows a bar chart of the Kolmogorov-Smirnov statistic *K_a_*, the outlier detection criterion from the previous figure. The bars are shown in the original order of the arrays. Based on the distribution of the values across all arrays, a threshold of 0.647 was determined, which is indicated by the vertical line. None of the arrays exceeded the threshold and was considered an outlier.

**[- Figure 6: Density plots.](javascript:%20toggle('dens'))**

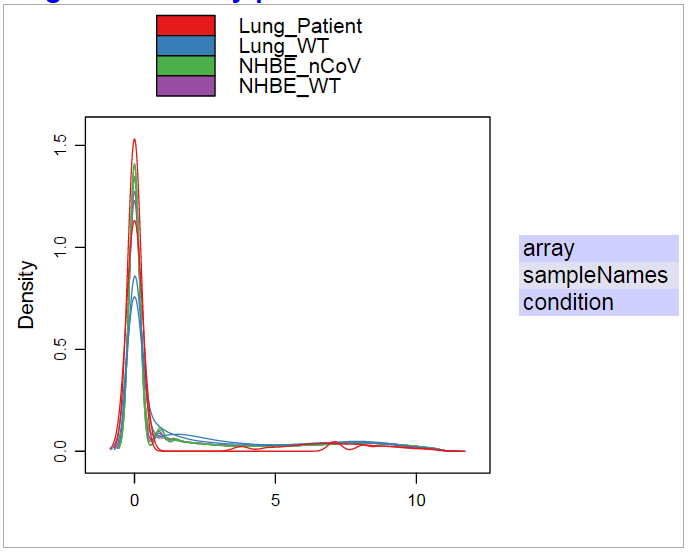

**Figure 6** shows density estimates (smoothed histograms) of the data. Typically, the distributions of the arrays should have similar shapes and ranges. Arrays whose distributions are very different from the others should be considered for possible problems. Various features of the distributions can be indicative of quality related phenomena. For instance, high levels of background will shift an array's distribution to the right. Lack of signal diminishes its right right tail. A bulge at the upper end of the intensity range often indicates signal saturation.

**Section 3: Variance mean dependence**

**[- Figure 7: Standard deviation versus rank of the mean.](javascript:%20toggle('msd'))**

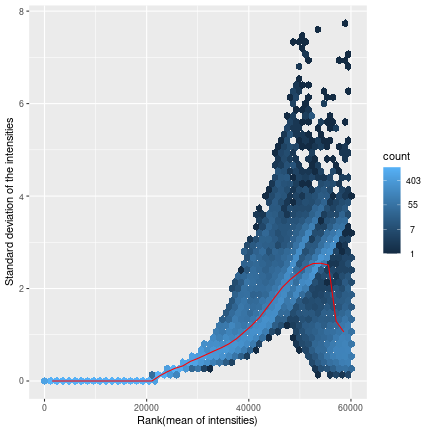

**Figure 7** shows a density plot of the standard deviation of the intensities across arrays on the *y*-axis versus the rank of their mean on the *x*-axis. The red dots, connected by lines, show the running median of the standard deviation. After normalisation and transformation to a logarithm (-like) scale, one typically expects the red line to be approximately horizontal, that is, show no substantial trend. In some cases, a hump on the right hand of the x-axis can be observed and is symptomatic of a saturation of the intensities.

**Section 4: Individual array quality**

**[- Figure 8: MA plots.](javascript:%20toggle('ma'))**

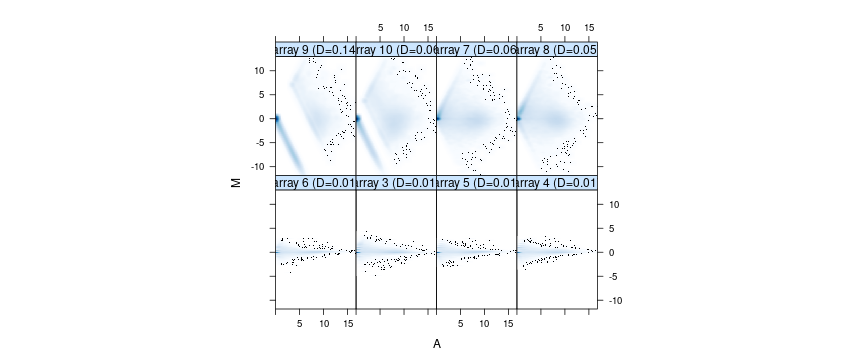

**Figure 8** shows MA plots. M and A are defined as:

M = log_2_(I_1_) - log_2_(I_2_)

A = 1/2 (log_2_(I_1_)+log_2_(I_2_)),

where I_1_ is the intensity of the array studied,and I_2_ is the intensity of a "pseudo"-array that consists of the median across arrays. Typically, we expect the mass of the distribution in an MA plot to be concentrated along the M = 0 axis, and there should be no trend in M as a function of A. If there is a trend in the lower range of A, this often indicates that the arrays have different background intensities; this may be addressed by background correction. A trend in the upper range of A can indicate saturation of the measurements; in mild cases, this may be addressed by non-linear normalisation (e.g. quantile normalisation).
Outlier detection was performed by computing Hoeffding's statistic *D_a_* on the joint distribution of A and M for each array. Shown are first the 4 arrays with the highest values of *D_a_*, then the 4 arrays with the lowest values. The value of *D_a_* is shown in the panel headings. 0 arrays had *D_a_*>0.15 and were marked as outliers. For more information on Hoeffing's *D*-statistic, please see the manual page of the function hoeffd in the Hmisc package.

**[- Figure 9: Outlier detection for MA plots.](javascript:%20toggle('out%20ma'))**

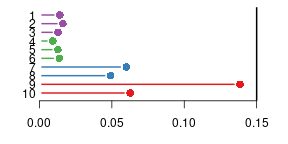

**Figure 9** shows a bar chart of the *D_a_*, the outlier detection criterion from the previous figure. The bars are shown in the original order of the arrays. A threshold of 0.15 was used, which is indicated by the vertical line. None of the arrays exceeded the threshold and was considered an outlier.
